# Lysosome-Dependent Sphingolipid Regulation as a potential therapeutic Target for Cohen Syndrome

**DOI:** 10.1101/2025.09.04.674037

**Authors:** Fabrizio Vacca, Renuka Prasad, Huda Barakullah, Romain Da Costa, Stefania Vossio, Dimitri Moreau, Woong Sun, Howard Riezman, Muhammad Ansar

**Author notes:** These authors share last authorship (Howard Riezman, Muhammad Ansar).

## Abstract

Cohen Syndrome (CS) is a rare autosomal recessive disorder caused by biallelic mutations in the *VPS13B* gene, affecting approximately 50,000 individuals worldwide. Clinical features include postnatal microcephaly, developmental delay, intellectual disability, neutropenia, and retinal dystrophy. VPS13B belongs to the bridge-like lipid transfer protein (BLTP) family, which also includes VPS13A, VPS13C, and VPS13D in mammals. Although its precise function remains unclear, VPS13B localizes to the Golgi complex, and its loss leads to Golgi fragmentation, a consistent cellular phenotype observed in VPS13B-deficient models. We used the rescue of this cell-autonomous phenotype as the basis for a microscopy-based high-throughput screening assay, through which we identified several small molecules capable of restoring Golgi morphology. Most of these compounds shared a common mechanism of action, relying on lipid accumulation in acidic organelles due to their cationic amphiphilic properties (CADs). Lipidomic profiling revealed a reduction in C18-N-acyl sphingolipids as a characteristic feature of VPS13B knockout (KO) cells, a defect that was reversed by the majority of the identified compounds. To evaluate the physiological relevance of these findings, we tested two compounds, azelastine and raloxifene, in cortical organoids (COs) derived from VPS13B KO human pluripotent stem cells. These organoids exhibited smaller size and reduced neurite outgrowth, reminiscent of the secondary microcephaly observed in CS patients. Treatment with either compound significantly recovered the neurite outgrowth phenotype, reinforcing physiological relevance of the compound effect. Taken together, our findings highlight a potential effect of the CAD on lysosome-dependent sphingolipid regulation, allowing the recovery of Golgi integrity and partial rescue in the cortical organoid CS model. Although additional studies are required to delineate the exact molecular targets, this work uncovers a potential mechanism that could be leveraged for the treatment of CS.

## INTRODUCTION

Cohen Syndrome (CS) is a rare genetic disease caused by biallelic mutations in the *VPS13B* gene, affecting an estimated 50’000 people worldwide^1^. It is a multisystemic disorder and main common symptoms include postnatal microcephaly, developmental delay, intellectual disability, facial dysmorphism, neutropenia, hypotonia and loss of vision due to progressive retinal dystrophy^2^; ^3^. *VPS13B*, along with *VPS13A, VPS13C and VPS13D*, is homologous of the yeast gene *VPS13* and belongs to a recently characterized protein family known as bridge-like lipid transfer protein (BLTP)^4^; ^5^. These proteins are organized as a long hydrophobic tunnel composed of repeated units named RBG (repeated beta-groove)^6–8^ and are thought to mediate lipid transfer through the hydrophobic tunnel between adjacent organelle membranes at membrane-contact sites (MCS)^9^; ^10^. Proteins of the BLTP family have diverse sub-cellular localizations and are often found at MCS connecting the ER with other organelles (see Hanna et al for a review^5^). VPS13B is instead localized to the Golgi complex and the more striking cellular phenotype observed upon VPS13B loss is fragmentation of this organelle^11^. Although the precise role of VPS13B in Golgi structure maintenance remains unclear, recent studies have provided insights into its mechanistic function. VPS13B has been implicated in connecting cis-and trans-Golgi cisternae in co-operation with its binding partner FAM177A1^12^ as well as in the formation of tubular ERGIC structures at the ER-Golgi interface by interacting with Sec23IP protein^13^. Moreover, silencing of VPS13B in hippocampal neuron cultures impairs neurite outgrowth and disrupts Golgi alignment with axonal directional growth, indicating a critical role in neuronal polarization during development^14^. Given the recognized importance of the Golgi complex in cell polarization-mediated processes during neurodevelopment^15^; ^16^ and the clinical similarities between CS and other disorders caused by mutations in genes essential for Golgi morphology^17^, it is reasonable to hypothesize that Golgi dysfunction plays a significant role in CS pathology.

Despite recent progress in elucidating VPS13B function, no therapy is currently available to prevent or mitigate central nervous system (CNS)-associated symptoms in CS patients, in contrast to metabolic and immunological manifestations, which are better managed in clinical practice^18^; ^19^. The limited understanding of the pathological mechanism at the molecular level also hampers target-based drug discovery efforts. In this study, we employed a phenotypic screening approach based on high-content microscopy to analyze Golgi morphology recovery in *VPS13B KO* cells. We screened the Prestwick Chemical Library (PCL, 1,280 compounds) and identified several small molecules capable of restoring Golgi structure. As a complementary approach, we performed a lipid composition analysis of *VPS13B* KO cells to elucidate specific roles of the protein in intracellular lipid metabolic processes and identify potential therapeutic targets.

## RESULTS

### Identification of compounds restoring Golgi morphology by high-content screening

As a cellular model for high-throughput screening for small molecules correcting CS phenotype, we generated VPS13B KO HeLa cell clones by CRISPR-mediated insertion/deletion (INDEL) mutations. We selected multiple clones raised with two different sgRNAs and confirmed the presence of homozygous or compound heterozygous frameshift mutations by genomic sequencing (Fig. S1A). VPS13B complete depletion at the protein level was confirmed by western blot on clone 2D9, that was finally selected to perform the compound screening (Fig. S1B). In agreement with previous observations in different cell types^11^; ^14^; ^19^, all selected clones present a clear fragmentation of the Golgi complex evidenced by staining with the cis-Golgi protein GM130 excluding a clone-dependent effect for this phenotype (Fig. S2). Clone 2D9 (VPS13B KO) and parental HeLa cells were used to set up quantitative analysis of Golgi morphology and assessing screening robustness and dynamic range. Cells stained with either GM130 or the trans-Golgi network (TGN) protein GOLPH3, display a similar fragmentation phenotype in VPS13B KO cells (Fig. 1A). For image analysis of Golgi morphology, after cell segmentation, multiple parameters were extracted for single cells, describing size, number, shape and cellular distribution of Golgi fragments (Fig. 1B). Golgi fragment average area and perimeter, number of Golgi fragments per cell, and shape factor average present a very good separation of WT from VPS13B KO cells with both GM130 (Fig. 1C) and GOLPH3 (Fig. S3A) as markers. GM130 with z-factor (Z’)^20^ above 0.5 for each individual parameter was selected as screening readout. Similar results were obtained in other VPS13B KO clones (Fig S3B, C). We confirmed that the fragmentation of Golgi complex is fully recovered by over-expressing the WT protein in VPS13B KO cells (Fig 1D, E). In order to search for additional readouts, Intracellular markers of other organelles such as early (EEA1) and late (LAMP1, Bis(monoacylglycero)phosphate/Lysobisphosphatidic acid (BMP/LBPA)) endocytic compartments and lipid droplets (PLIN2) were also tested: we could not detect however any significant difference between WT and VPS13B KO cells (Fig. S4). The screening was performed in duplicate on 384-well plates pre-spotted with compounds at a final concentration of 10 µM. Cells were fixed and stained 24 hours after plating. 32 wells of vehicle-treated WT and VPS13B KO cells were included in each plate for internal normalization. After image analysis and extraction of Golgi morphology descriptors, a recovery ranking of compound was established based on the ability of compound to minimize the 4-dimentional distance (considering fragment Golgi average area and perimeter, shape factor and number of fragments per cell. See Methods for details) to WT cells values. The separation between WT and KO untreated control was very robust in the whole screening with compounds showing a continuous degree range of correction levels (from 0 to 1) (Fig. 2A, B). An arbitrary threshold was set at 0.4 (including a total of 50 compounds) to define positive hits for further analysis (Table 1, full ranking in File S1). This value reflects the visual assessment of Golgi morphology recovery after visual inspection of the images (Fig. 2C). One single compound was excluded from the ranking due to evidence of analysis artifacts associated with severely altered cell morphology. Most of the molecules composing the PCL are approved drugs with defined protein targets. This allows a first scrutiny of potential mechanism of action (MOA): The hits have a heterogeneous group of targets with D1/D2 dopamine receptors, Serotonin and norepinephrine transporters and Histamine H1 receptor being the more represented. These compounds are labeled as in Psycholeptics, Psychoanaleptics and Antihistamines in level 2 Anatomical Therapeutic Chemical (ATC) classification system (Fig. 2D). From a chemical perspective, half of them (26/50) are aromatic tricyclic compounds (either homo-or hetero-cyclic). Most of the remaining compounds are also aromatic and polycyclic, with the exception of two corticosteroids, budesonide and ciclesonide. Moreover, the vast majority of the hits (47/50) have secondary or tertiary amine groups with basic pK_a_ >7, which, together with elevated hydrophobicity, estimated by AlogP value^21^ (logP>3.5), categorize them as lysosomotrophic compounds (Fig. 2E). These molecules, also known as cationic amphiphilic drugs (CAD), get trapped in the lumen of acidic organelles after protonation of their amine group, causing an impairment of lipase activity and lysosomal lipid storage^22–24^. CAD, with a hit frequency of 0.9 are indeed very significantly enriched with respect to their overall frequency in the PCL (0.16) (Fig 2F).

**Figure 1.**
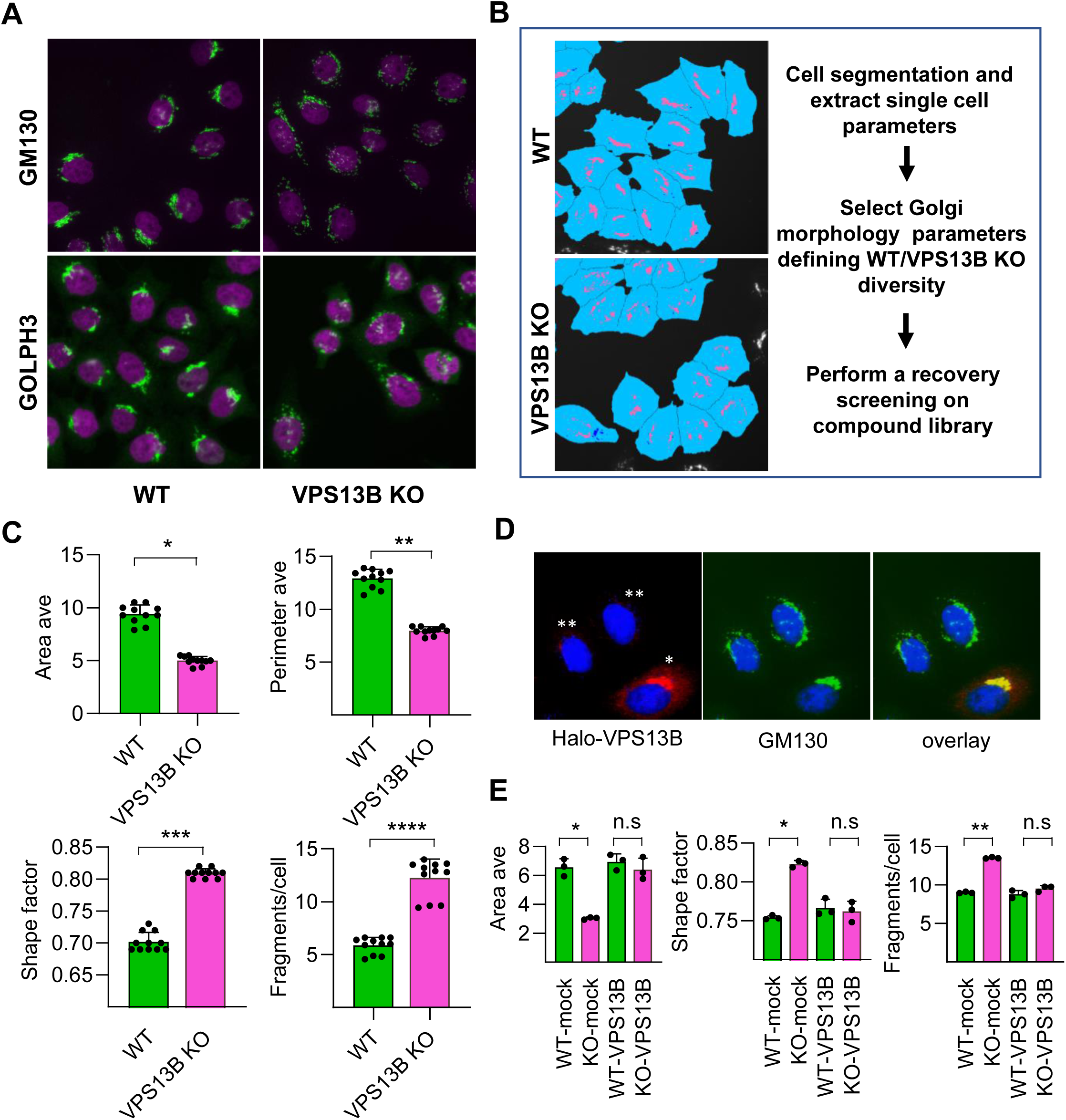
Definition of Golgi morphology as a readout of high-content screening. **A)** Representative images of fixed WT and VPS13B KO HeLa cells stained for GM130 (upper panel, green) and GOLPH3 (lower panel, green). Nuclei stained with Hoechst 33342 (magenta). **B)** simplified workflow of image analysis performed in the screening with an example of cell segmentation (light blue) with a Golgi mask (rose) based on GM130 staining. **C)** Analysis of Golgi morphology parameters extracted from GM130 images. Average area and perimeter of Golgi fragments (Area Ave, Perimeter Ave); Shape factor average; average number of Golgi fragments per cell. Analysis based on 1000-1500 cells/point (*p<0.0001, z’=0.52; **p<0.0001, z’=0.64; ***p<0.0001, z’=0.83; ****p<0.0001, z’=0.66; paired t-test, 2-tails; N=11) **D)** representative images of VPS13B KO cells transfected with Halo-VPS13B (red) and stained for GM130 (green). Nuclei stained with Hoechst 33342 (blue). *transfected, **non-transfected. **E)** Analysis of Golgi parameters (Average area, Shape factor, Fragments/cell) performed on mock or Halo-VPS13B transfected HeLa cells (WT and VPS13B KO). For Halo-VPS13B transfected cells analysis was performed only in Halo-positive cells (*p<0.01; **p<0.001; paired t-test, 2-tails; N=3)

**Figure 2.**
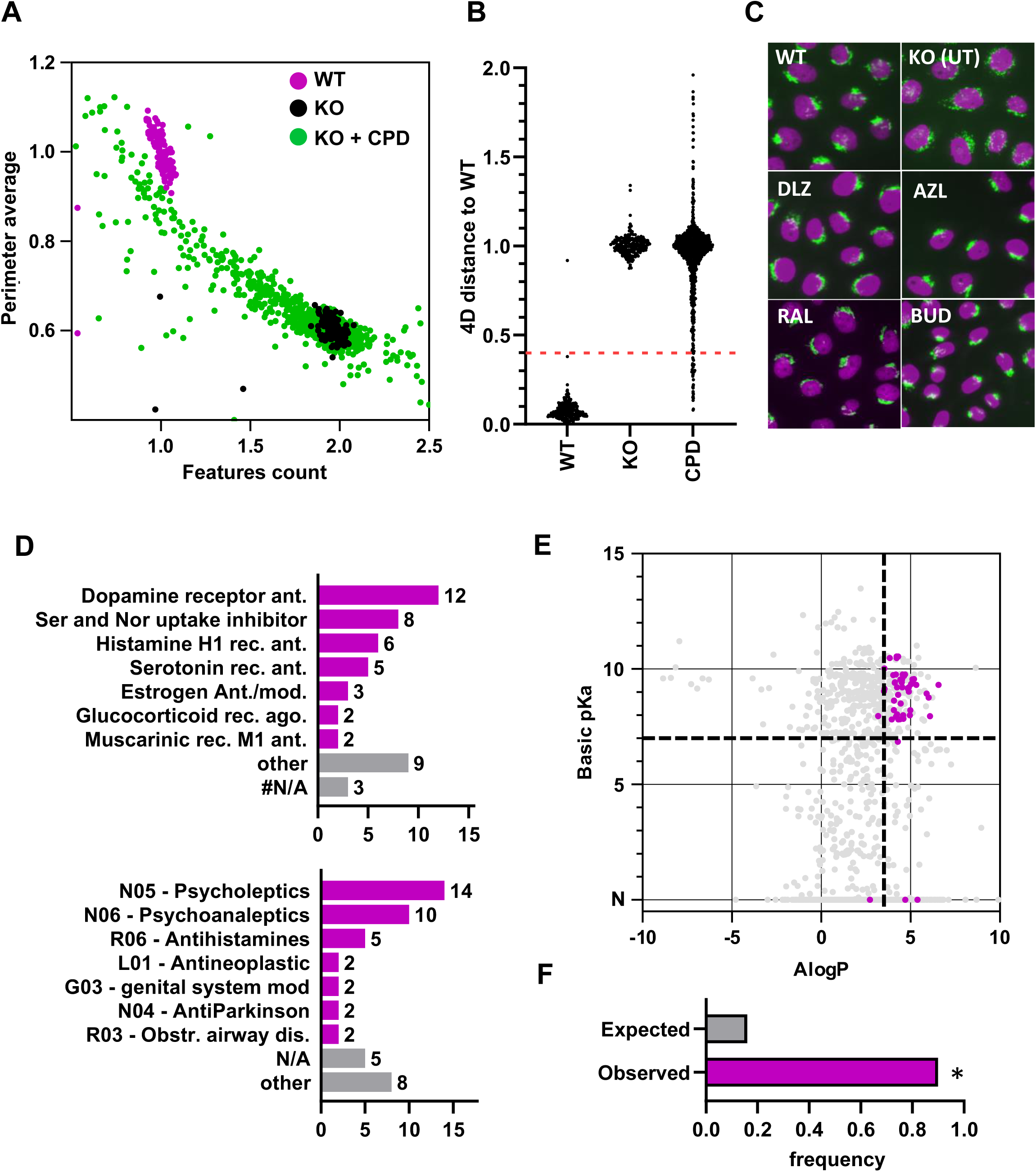
Phenotypic screening identifies compounds that restore Golgi morphology in VPS13B-deficient cells. **A)** Schematic 2D representation of screening output, plotting 2 parameters (Golgi perimeter average, Features count/cell) for WT (magenta) and untreated VPS13B KO (black) replicates and compound-treated VPS13B KO cells (green), each dot representing a different compound. Analysis based on 1000-1500 cells/point (N=2). **B)** Representation of the data as the distance to WT based on 4 parameters (Golgi fragment average area, Golgi fragment average perimeter, shape factor, Golgi fragments/cell) plotted for WT and untreated VPA13B KO cells and for individual compound-treated VPS13B KO cells (CPD). (N=2). **C)** Representative screening images stained for GM130 (green) and Hoechst 33342 (magenta) showing WT and VPS13B KO HeLa cells and 4 different compound-treated cells: dilazep (DLZ); azelastine (AZL); raloxifene (ral); budesonide (BUD), 10 µM. **D)** Distribution of top 50 screening hits by primary therapeutic protein target (upper panel) and 2nd level ATC classification group (lower panel). Data obtained from PubChem (pubchem.ncbi.nlm.nih.gov). **E)** Screening library compound plotted according to predicted basic pKa and AlogP value. Top 50 hits are shown in magenta. Data obtained from ChEMBL (ebi.ac.uk/chembl). **F)** Expected and observed hit frequency of CAD compounds in the library. *p<0.0001, Wilson-Brown Binomial test.

**Table 1.**
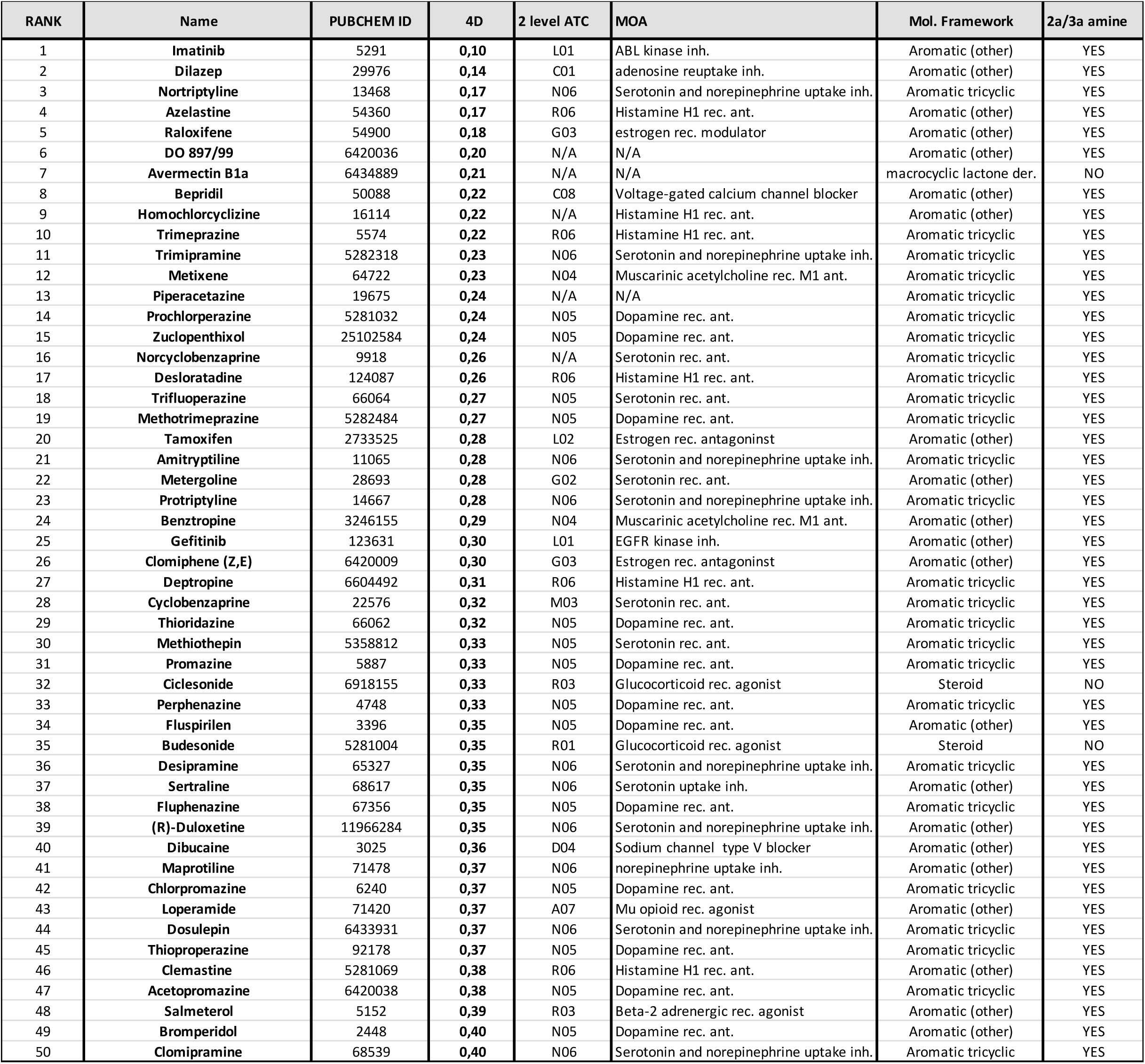
List of the top 50 compounds ranked on the basis of the 4-dimensional image parameter (4D) for the recovery of Golgi morphology in VPS13B KO cells.

| RANK | Name | PUBCHEM ID | 4D | 2 level ATC | MOA | Mol. Framework | 2a/3a amine |
| --- | --- | --- | --- | --- | --- | --- | --- |
| 1 | Imatinib | 5291 | 0,10 | L01 | ABL kinase inh. | Aromatic (other) | YES |
| 2 | Dilazep | 29976 | 0,14 | C01 | adenosine reuptake inh. | Aromatic (other) | YES |
| 3 | Nortriptyline | 13468 | 0,17 | N06 | Serotonin and norepinephrine uptake inh. | Aromatic tricyclic | YES |
| 4 | Azelastine | 54360 | 0,17 | R06 | Histamine H1 rec. ant. | Aromatic (other) | YES |
| 5 | Raloxifene | 54900 | 0,18 | G03 | estrogen rec. modulator | Aromatic (other) | YES |
| 6 | DO 897/99 | 6420036 | 0,20 | N/A | N/A | Aromatic (other) | YES |
| 7 | Avermectin B1a | 6434889 | 0,21 | N/A | N/A | macrocyclic lactone der. | NO |
| 8 | Bepidil | 50088 | 0,22 | C08 | Voltage-gated calcium channel blocker | Aromatic (other) | YES |
| 9 | Homochlorcyclizine | 16114 | 0,22 | N/A | Histamine H1 rec. ant. | Aromatic (other) | YES |
| 10 | Trimeprazine | 5574 | 0,22 | R06 | Histamine H1 rec. ant. | Aromatic tricyclic | YES |
| 11 | Trimipramine | 5282318 | 0,23 | N06 | Serotonin and norepinephrine uptake inh. | Aromatic tricyclic | YES |
| 12 | Metixene | 64722 | 0,23 | N04 | Muscarinic acetylcholine rec. M1 ant. | Aromatic tricyclic | YES |
| 13 | Piperacetazine | 19675 | 0,24 | N/A | N/A | Aromatic tricyclic | YES |
| 14 | Prochlorperazine | 5281032 | 0,24 | N05 | Dopamine rec. ant. | Aromatic tricyclic | YES |
| 15 | Zuclopenthixol | 25102584 | 0,24 | N05 | Dopamine rec. ant. | Aromatic tricyclic | YES |
| 16 | Norcyclobenzaprine | 9918 | 0,26 | N/A | Serotonin rec. ant. | Aromatic tricyclic | YES |
| 17 | Desloratadine | 124087 | 0,26 | R06 | Histamine H1 rec. ant. | Aromatic tricyclic | YES |
| 18 | Trifluoperazine | 66064 | 0,27 | N05 | Serotonin rec. ant. | Aromatic tricyclic | YES |
| 19 | Methotrimeprazine | 5282484 | 0,27 | N05 | Dopamine rec. ant. | Aromatic tricyclic | YES |
| 20 | Tamoxifen | 2733525 | 0,28 | L02 | Estrogen rec. antagonist | Aromatic (other) | YES |
| 21 | Amityptiline | 11065 | 0,28 | N06 | Serotonin and norepinephrine uptake inh. | Aromatic tricyclic | YES |
| 22 | Metergoline | 28693 | 0,28 | G02 | Serotonin rec. ant. | Aromatic (other) | YES |
| 23 | Protriptyline | 14667 | 0,28 | N06 | Serotonin and norepinephrine uptake inh. | Aromatic tricyclic | YES |
| 24 | Benztropine | 3246155 | 0,29 | N04 | Muscarinic acetylcholine rec. M1 ant. | Aromatic (other) | YES |
| 25 | Gefitinib | 123631 | 0,30 | L01 | EGFR kinase inh. | Aromatic (other) | YES |
| 26 | Clomiphene (Z,E) | 6420009 | 0,30 | G03 | Estrogen rec. antagonist | Aromatic (other) | YES |
| 27 | Deptropine | 6604492 | 0,31 | R06 | Histamine H1 rec. ant. | Aromatic tricyclic | YES |
| 28 | Cyclobenzaprine | 22576 | 0,32 | M03 | Serotonin rec. ant. | Aromatic tricyclic | YES |
| 29 | Thioridazine | 66062 | 0,32 | N05 | Dopamine rec. ant. | Aromatic tricyclic | YES |
| 30 | Methiothepin | 5358812 | 0,33 | N05 | Serotonin rec. ant. | Aromatic tricyclic | YES |
| 31 | Promazine | 5887 | 0,33 | N05 | Dopamine rec. ant. | Aromatic tricyclic | YES |
| 32 | Ciclesonide | 6918155 | 0,33 | R03 | Glucocorticoid rec. agonist | Steroid | NO |
| 33 | Perphenazine | 4748 | 0,33 | N05 | Dopamine rec. ant. | Aromatic tricyclic | YES |
| 34 | Fluspirilen | 3396 | 0,35 | N05 | Dopamine rec. ant. | Aromatic (other) | YES |
| 35 | Budesonide | 5281004 | 0,35 | R01 | Glucocorticoid rec. agonist | Steroid | NO |
| 36 | Desipramine | 65327 | 0,35 | N06 | Serotonin and norepinephrine uptake inh. | Aromatic tricyclic | YES |
| 37 | Sertraline | 68617 | 0,35 | N06 | Serotonin uptake inh. | Aromatic (other) | YES |
| 38 | Fluphenazine | 67356 | 0,35 | N05 | Dopamine rec. ant. | Aromatic tricyclic | YES |
| 39 | (R)-Duloxetine | 11966284 | 0,35 | N06 | Serotonin and norepinephrine uptake inh. | Aromatic (other) | YES |
| 40 | Dibucaine | 3025 | 0,36 | D04 | Sodium channel type V blocker | Aromatic (other) | YES |
| 41 | Maprotiline | 71478 | 0,37 | N06 | norepinephrine uptake inh. | Aromatic (other) | YES |
| 42 | Chlorpromazine | 6240 | 0,37 | N05 | Dopamine rec. ant. | Aromatic tricyclic | YES |
| 43 | Loperamide | 71420 | 0,37 | A07 | Mu opioid rec. agonist | Aromatic (other) | YES |
| 44 | Dosulepin | 6433931 | 0,37 | N06 | Serotonin and norepinephrine uptake inh. | Aromatic tricyclic | YES |
| 45 | Thiopropazine | 92178 | 0,37 | N05 | Dopamine rec. ant. | Aromatic tricyclic | YES |
| 46 | Clemastine | 5281069 | 0,38 | R06 | Histamine H1 rec. ant. | Aromatic (other) | YES |
| 47 | Acetopromazine | 6420038 | 0,38 | N05 | Dopamine rec. ant. | Aromatic tricyclic | YES |
| 48 | Salmeterol | 5152 | 0,39 | R03 | Beta-2 adrenergic rec. agonist | Aromatic (other) | YES |
| 49 | Bromperidol | 2448 | 0,40 | N05 | Dopamine rec. ant. | Aromatic (other) | YES |
| 50 | Clomipramine | 68539 | 0,40 | N06 | Serotonin and norepinephrine uptake inh. | Aromatic tricyclic | YES |

### Lysosomal lipid storage mediates the recovery of Golgi morphology

Given the predicted lysosomotrophic properties of most of the hits, we tested the ability of selected compounds to induce lysosomal lipid storage and analyzed the link with recovery of Golgi morphology. The composition of the material accumulating in endo/lysosomal organelles can be very heterogeneous, but the lipid storage phenotype can be easily monitored by immuno-fluorescence (IF) by staining BMP/LBPA which is known to accumulate also in genetic lysosomal storage diseases (LSD) such as Niemann-Pick C (NPC)^25^; ^26^ or with filipin dye which labels unesterified cholesterol^27^; ^28^. As shown in figure 3A, B, four different compounds (azelastine, dilazep, trifluoperazine and raloxifene) cause a dose-dependent increase of LBPA and cholesterol staining in the micromolar range. As a comparison, we treated cells with U18666A which causes lysosomal lipid storage by direct inhibition of Niemann-Pick C1 (NPC1) protein function^29^. Interestingly, cell treatment with U18666A also partially correct Golgi morphology (Fig. S5) The selected compounds also cause a strongly increased staining with LipidTOX phospholipidosis detection reagent (Fig. 3C), a lipophilic fluorescent dye previously used in drug screenings to detect drug-induced lipid storage, normally considered as a potentially deleterious side effect^22^;^30^; ^31^. Even if the term phospholipidosis is commonly associated with lysosomal lipid storage, we observed that the LipidTOX dye mostly accumulates in Transferrin receptor (TFRC) containing compartments, with only a small portion in LAMP1-positive late endosomes/lysosomes and no detectable co-localization with the lipid droplet marker PLIN2 (Fig. S6). While we still assume that most lipid storage takes place in late endosomal compartments, this observation likely indicates a broader traffic impairment of the endocytic system after drug treatment, not limited to late endosomes/lysosomes. It is also worth noticing that, when testing the effect of selected drugs (azelastine, dilazep and raloxifene) in WT cells, we observe minor but measurable changes in Golgi morphology (Fig. S7). Although these changes were substantially less pronounced than those observed in VPS13B KO cells and were barely detectable by visual inspection, they nonetheless indicate that the effects of these drugs on Golgi organization are more general and not restricted to the KO context.

In order to assess a causative role of lysosomal storage in Golgi phenotypic recovery we silenced either Acid sphingomyelinase (SMPD1) or NPC1, whose loss-of-function mutations cause Niemann-Pick A/B and C diseases, repectively^32^. NPC1 knock-down (KD) causes a complete recovery of Golgi phenotype, while SMPD1 KD partially rescued normal Golgi morphology (Fig. 3D, E). These findings mirror the effects of lysosomotropic compounds, suggesting that the observed Golgi recovery under CAD treatment is indeed mediated by lipid accumulation in endosomes and lysosomes in VPS13B KO cells.

**Figure 3.**
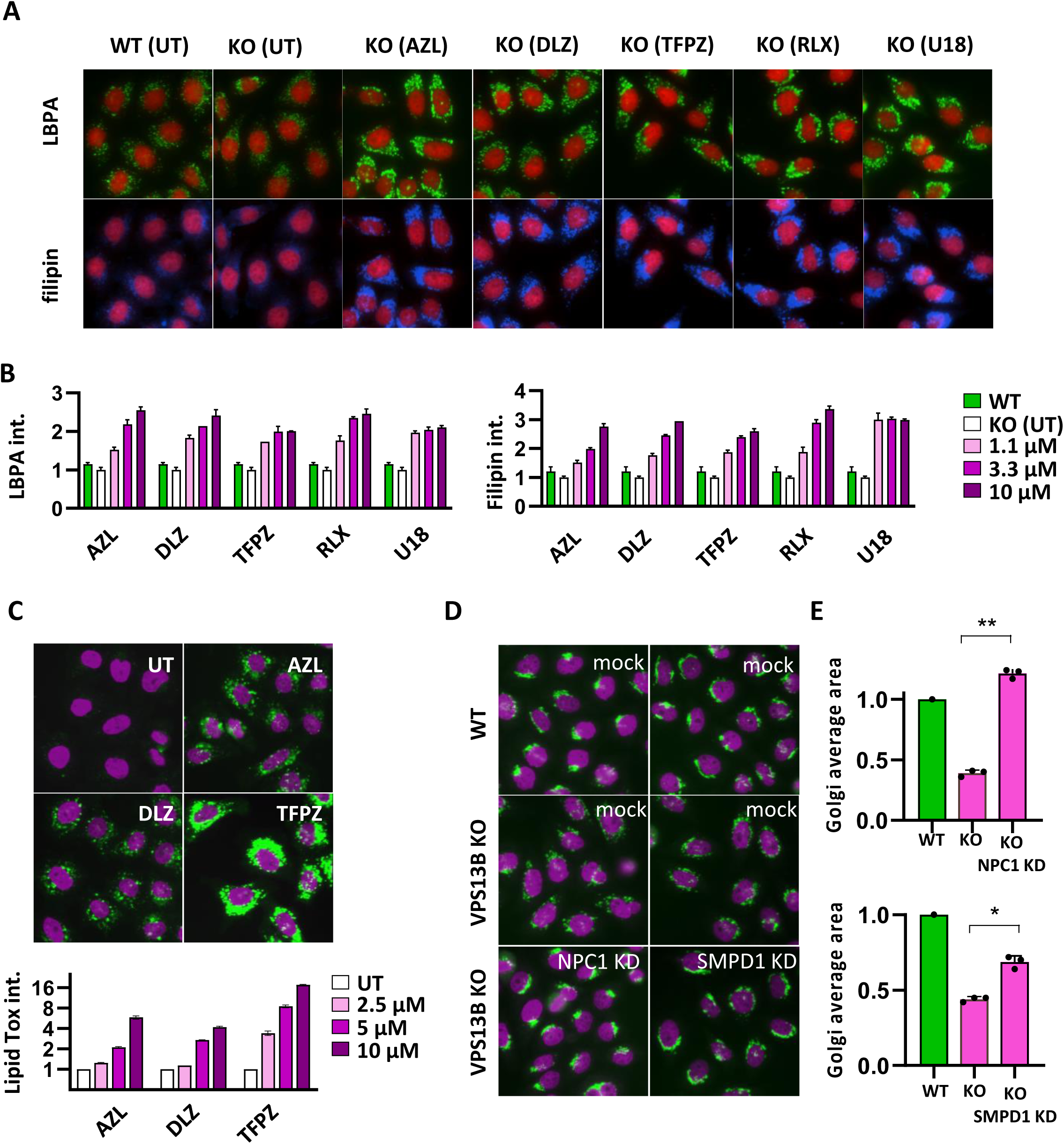
Lysosomal lipid storage mediates the recovery of Golgi morphology. **A)** Representative images of WT Hela cells or VPS13B KO cells treated with: azelastine (AZL); dilazep (DLZ); trifluoperazine (TFPZ); raloxifene (RLX); U18666A (U18). 10 µM, 24 h. cells were fixed and stained with an anti-LBPA antibody (upper panel) of with Filipin (lower panel). **B)** Integrated intensity/cell quantification of LBPA and Filipin staining at different compound doses (1.1, 3.3, 10 µM) for 24h. Relative to untreated (a.u). Analysis based on 1000-1500 cells/point (N=2). **C)** Representative images of LipidTox accumulation in cells treated with azelastine (AZL); dilazep (DLZ); trifluoperazine (TFPZ). 10 µM, 24 h (upper panel). Integrated intensity/cell quantification of LipidTox at different compound doses (2.5, 5, 10 µM) for 24h. Relative to untreated (a.u). Analysis based on 1000-1500 cells/point (N=2). **D)** Representative images of WT or VPS13B KO transfected with non-targeting (mock) or targeting NPC1 or SMPD1 for 72 hours. Cells were stained for GM130 (green) and Hoechst 33342 (magenta). **E)** Analysis of Golgi average area (value relative to WT cells) in NPC1 and SMPD1 KD. Analysis based on 1000-1500 cells/point. (*p<0.001, **p<0.0001. paired t-test, 2-tails; N=3)

We performed dose-response curves (DRC) (from 78 nM to 10 µM) for 14 compounds selected among the top 50 in the ranking. The upper concentration limit was kept to 10 µM because we were mostly motivated to explore compounds with better potency, rather that maximal efficacy at high doses. The compounds were picked in order to represent the structural diversity among the hits. As an additional criterium, finalized to potential future *in vivo* applications, compounds with lower reported toxicity were preferred. Since most of the compounds are predicted to be lysosomotrophic (with the notable exception of the glucocorticoids budesonide and ciclesonide), we included LipidTOX staining in the DRC in order to correlate lipid storage and recovery of Golgi phenotype. As can be clearly observed in DRC charts (Fig. 4A-D; Fig. S8), for all CAD, the concentration at which the effect on Golgi phenotype start to be measurable falls in the low micromolar range and parallels the rise in lipid storage measured with LipidTOX. This observation reinforces the link between lipid storage and Golgi morphological recovery. Since most of the compounds did not reach a plateau level within the 10 µM limit, it was not possible to define a proper EC_50_ value for the response. As an estimate of the potency, we report here the value at which a 50% recovery of Golgi morphology is reached according to the same parameters used to rank the compounds in the primary screening and the maximal efficacy at 10 µM (Table 2; File S2). Although all predicted lysosomotropic drugs were confirmed by exhibiting LipidTOX storage, the extent of accumulation varied significantly, ranging from a 5-fold increase with Bepridil to a 30-fold increase with trifluoperazine and sertraline (Table 2). While there is some correlation between LipidTOX storage and potency, certain highly potent compounds, such as dilazep and azelastine, do not induce the strongest LipidTOX accumulation (8-and 12-fold increases, respectively). Budesonide requires separate consideration. As a glucocorticoid, it has been confirmed not to be lysosomotropic (Fig. 4D). It exhibits slightly higher potency than the other compounds (2 µM) and maximal efficacy at 5 µM, followed by a decline at 10 µM. A closer examination of the phenotype suggests that this reduction in efficacy is not due to increased Golgi fragmentation but rather to excessive Golgi clustering compared to the wild-type (WT) condition (Fig. S9).

**Figure 4.**
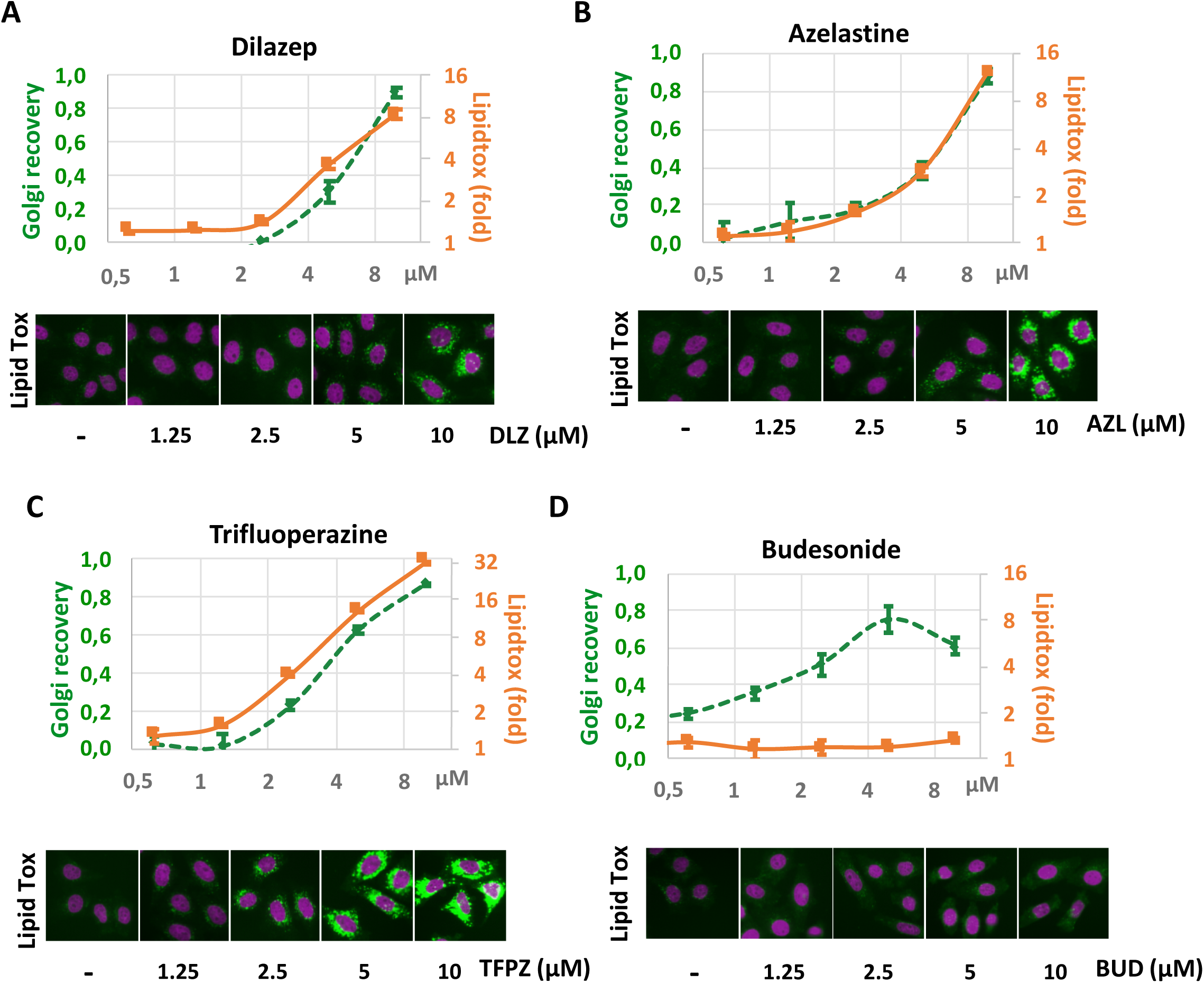
Dose response curves of selected hits. **(A)** dilazep, **(B)** azelastine, **(C)** trifluoperazine, **(D) (upper panels)** displaying the Golgi recovery, expressed as the inverse (1-D) of 4-parametric distance to WT (green line) and LipidTox integrated intensity (relative to WT, a.u., orange line) Analysis based on 1000-1500 cells/point (N=2). **(lower panels)** Representative images of cells stained with LipidTox with 1.25 µM to 10 µM compound treatments.

**Table 2.** Data resume from dose-response curves. . In column 1, the values indicate the concentration (µM) at which a 50% recovery of Golgi morphology is achieved (based on the 4D value in WT cells). Column 2 indicates the max. recovery relative to WT (efficacy). Column 3 indicates the intensity of LipidTOX staining at maximal concentration (10 µM) relative to untreated cells

| Name | 50% recovery<br>(μM) | Max recovery<br>(1-4D) | LipidTOX at 10 μM |
| --- | --- | --- | --- |
| Budesonide | 2,10 | 0,75 | 1,3 |
| Trifluoperazine | 4,01 | 0,87 | 32,5 |
| Sertraline | 4,36 | 0,75 | 29,8 |
| Thioridazine | 5,25 | 0,66 | 23,0 |
| Nortriptyline | 5,29 | 0,80 | 15,1 |
| Azelastine | 6,21 | 0,88 | 12,1 |
| Dilazep | 6,6 | 0,90 | 8,4 |
| Chlorpromazine | 6,67 | 0,54 | 11,0 |
| Bepidil | 6,97 | 0,66 | 5,3 |
| (R)-Duloxetine | 7,90 | 0,67 | 12,8 |
| Desipramine | 8,47 | 0,65 | 10,2 |
| Trimipramine | 8,48 | 0,69 | 8,7 |
| Cyclobenzaprine | 8,52 | 0,65 | 8,4 |
| Amitryptiline | 9,54 | 0,55 | 7,5 |

To determine whether the observed Golgi recovery was transient, we extended the treatment duration to 72 hours for two selected compounds. Under these conditions, azelastine maintained a sustained beneficial effect on Golgi integrity (Fig. S10). In contrast, raloxifene proved to be toxic upon prolonged exposure, leading to extensive cell death and therefore precluding further analysis on Golgi morphology (data not shown).

### CADs do not prevent brefeldin A-and nocodazole-induced Golgi fragmentation but prevent statin-induced fragmentation

Treatment of cells with fungal metabolite brefeldin A (BFA) blocks ER to Golgi transport through inhibition of ARF1, causing a reversible disassembly and dispersion of Golgi complex^33–35^. Unlike cis/medial Golgi enzymes which are re-localized to the ER, GM130 appears fragmented but still co-colocalizing with other Golgi markers^36^. Microtubule depolymerization also causes reversible Golgi fragmentation with formation of functional mini-stacks^37^; ^38^. In order to test how general is the clustering effect of CADs, we tested their ability to prevent Golgi fragmentation induced by BFA or nocodazole treatment or to accelerate recovery after washout of these compounds. WT HeLa cells were kept in the presence of azelastine for 18 hours before treatment with either BFA for 1 hour (Fig. 5A) or nocodazole for 2 hours (Fig. 5B) to induce extensive Golgi complex fragmentation. After washout, the Golgi morphology was monitored until complete recovery (4 hours), keeping azelastine in the medium during the entire recovery period. azelastine was neither able to prevent fragmentation or to accelerate recovery for both conditions. Similar results were obtained with dilazep in cells treated with BFA (Fig. S11).

**Figure 5.**
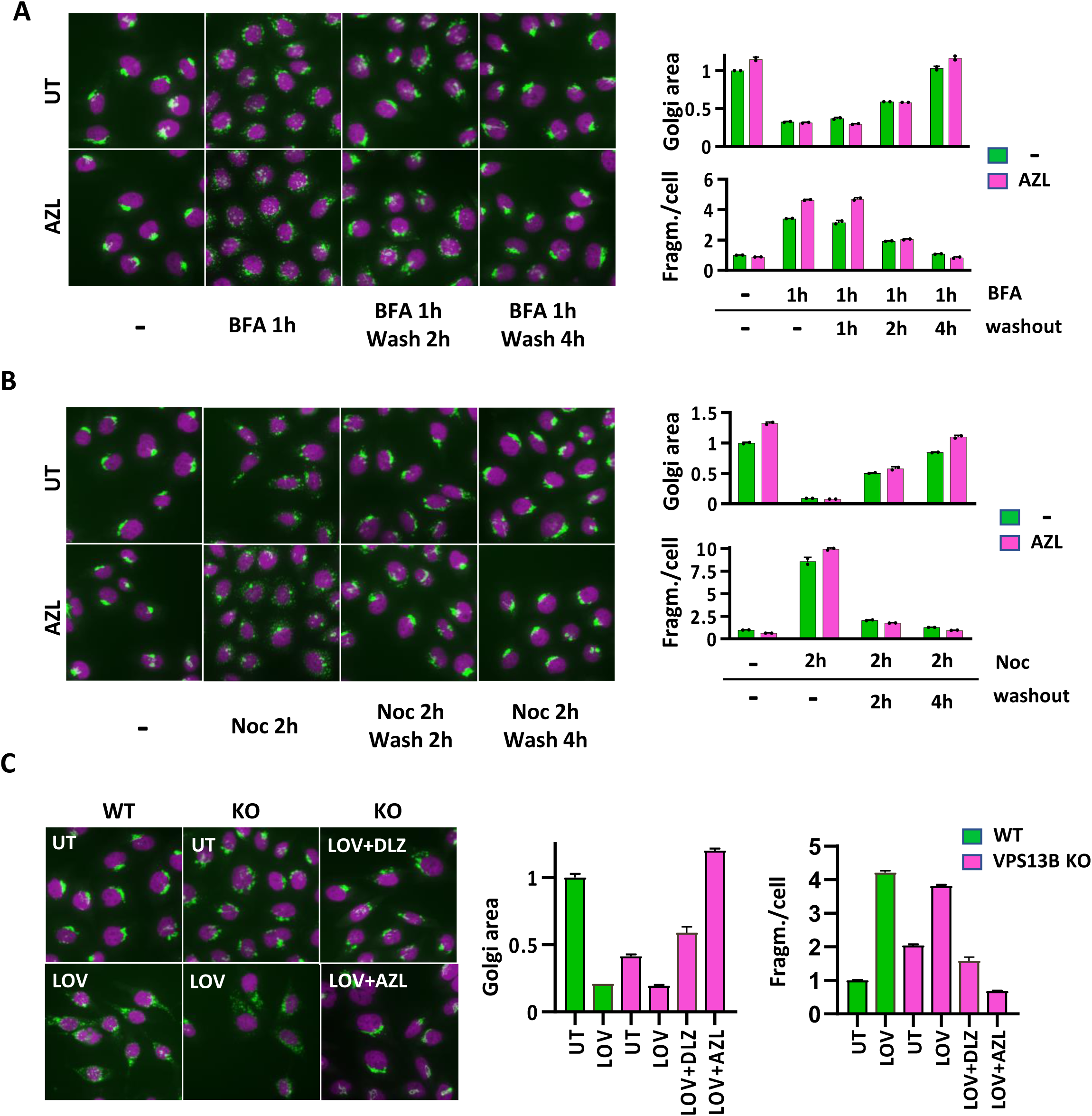
Differential effects of CADs on Golgi fragmentation induced by distinct pharmacological treatments. **A)** Representative images of WT HeLa cells treated (AZL) or not (UT) with azelastine 10µM for 24 hours. Cells were stained for GM130 (green) and Hoechst 33342 (magenta). Where indicated, cells have been subsequently with brefeldin A (BFA) 1 µg/ml for 1 hours and after 3 washes, let recover in full medium for 2 or 4 additional hours. In AZL samples, has been kept in all the course of the experiment. **B)** Representative images of WT HeLa cells treated (AZL) or not (UT) with azelastine 10µM for 24 hours. Cells were stained for GM130 (green) and Hoechst 33342 (magenta). Where indicated, cells have been subsequently with nocodazole (Noc) 5 µM for 2 hours and after 3 washes, let recover in full medium for 2 or 4 additional hours. In AZL samples, has been kept in all the course of the experiment. **(A, B right panels).** Graphs illustrating Golgi average area and average fragments/cell (values relative to UT cells) quantified on 1000-1500 cells/point (N=2). **C)** Representative images of WT or VPS13B KO HeLa cells treated (LOV) or not (UT) with lovastatin 5 µM for 24 hours. Cells were stained for GM130 (green) and Hoechst 33342 (magenta). Where indicated cells have been co-treated with either dilazep (DLZ) or azelastine (AZL) 10 µM. Graph illustrating Golgi average area and average fragments/cell (values relative to UT cells) quantified on 1000-1500 cells/point (N=2).

Although the screening was primarily focused on identifying compounds that restore Golgi morphology, we also observed several compounds that caused excessive fragmentation of the organelle.

Among those, we noticed the presence of 4 compounds belonging to the statin class (atorvastatin, lovastatin, fluvastatin, mevastatin) (File S1, Fig. S12). These molecules are inhibitors of HMG Co-A reductase, commonly used as cholesterol-lowering drugs. While we cannot exclude an effect due to cholesterol depletion, the more plausible explanation is that they act through the prenylation inhibition of Rab and SNARE proteins essential for Golgi morphology maintenance, as previously described^39^; ^40^. Whatever the mechanism of statin-induced Golgi dispersion, the concomitant treatment with dilazep and azelastine overcomes the effect re-establishing a compact Golgi in lovastatin-treated cells (Fig. 5C).

### Reduction of sphingolipids with C18 acyl chains in VPS13B KO cells

Since VPS13 is a *bona fide* lipid transfer protein, we performed lipidomic analysis in order to identify lipid alterations in VPS13B KO cells. Our goal was to gain insight into the mechanistic role of VPS13B in lipid metabolism and explore potential drug-mediated intervention strategies. Although the overall cellular lipid composition does not appear dramatically altered, analysis of lipid class abundance revealed specific changes. In VPS13B KO cells, we observed a 20–30% increase in lysophosphatidylcholines (LPC) and lysophosphatidylethanolamines (LPE), accompanied by a 10–20% decrease in sphingomyelin (SM) and trihexosylceramides (Hex3Cer) Meanwhile major glycerophospholipid (GP) species are not significantly altered (Fig. 6A, File S3). When analyzing individual lipid species, we noticed a decrease in the content of some specific SMs and ceramides (CER), notably SM 36:1, SM 36:2 and CER 36:1 as well as SM 38:1 and CER 38:1 (Fig. 6B). Given the assumption that sphingolipids in mammals are in vast majority built with an 18:1 sphingosine moiety^41^, these SM and ceramide species most likely contain C18 and C20 N-linked acyl chains, respectively. C18 SM is particularly intriguing as it has been shown to be highly enriched in COPI vesicles^42^ and to bind and regulate the function of TMED family proteins, which play crucial roles in Golgi-ER trafficking^43^; ^44^. Unlike tissues such as brain^45^ and skeletal muscle^46^ where they may represent the more abundant form of SM, C18 SM species constitute only a minor fraction of SM in HeLa cells (SM 36:1 <5%, SM 36:2 ∼1%). C18 ceramides are even less abundant. Nevertheless, even when expressed as a fraction of total SM, accounting for the overall SM reduction in VPS13B KO cells, C18 SM species are still significantly reduced (Fig. 6C, File S3). Some lipid species appear also to be increased in VPS13B KO cells. These include triacylglycerols (TG 58:1, TG 60:2, TG 60:3), phosphatidylcholines (PC 40:1), phosphatidylethanolamine (PE 42:1) as well as SM 42:1 and CER 44:1. With the notable exception of SM 42:1 representing ∼5% of total SM, these are very minor lipid species (below 1% of the relative lipid class, see File S3) and we were unable to find any reported specific function for these lipids. Since the observed lipid alterations could result from either a clonal effect or off-target effects of CRISPR-Cas9, we sought to validate our findings using an independent VPS13B KO clone (clone 3B8) raised with a different sgRNA as well as with a VPS13B WT clone (clone 3D10) isolated during the KO production process. As shown in figure 6D, clone 3B8 exhibits a reduction in SM 36:1, SM 36:1, CER 36:1 similarly to the clone used in the rest of the experiments and in the compound screening (clone 2D9). The clone 3D10 on the opposite does not display any significant difference from WT parental cells. Remarkably, a similar outcome was observed for other lipid species with altered levels in the KO clone 2D9 (Fig. S13). Conversely, the reduction in total SM was not observed in the 3B8 KO clone while there is an apparent decrease in Hes3cer levels was observed, though it did not reach statistical significance (Fig. S13).

**Figure 6.**
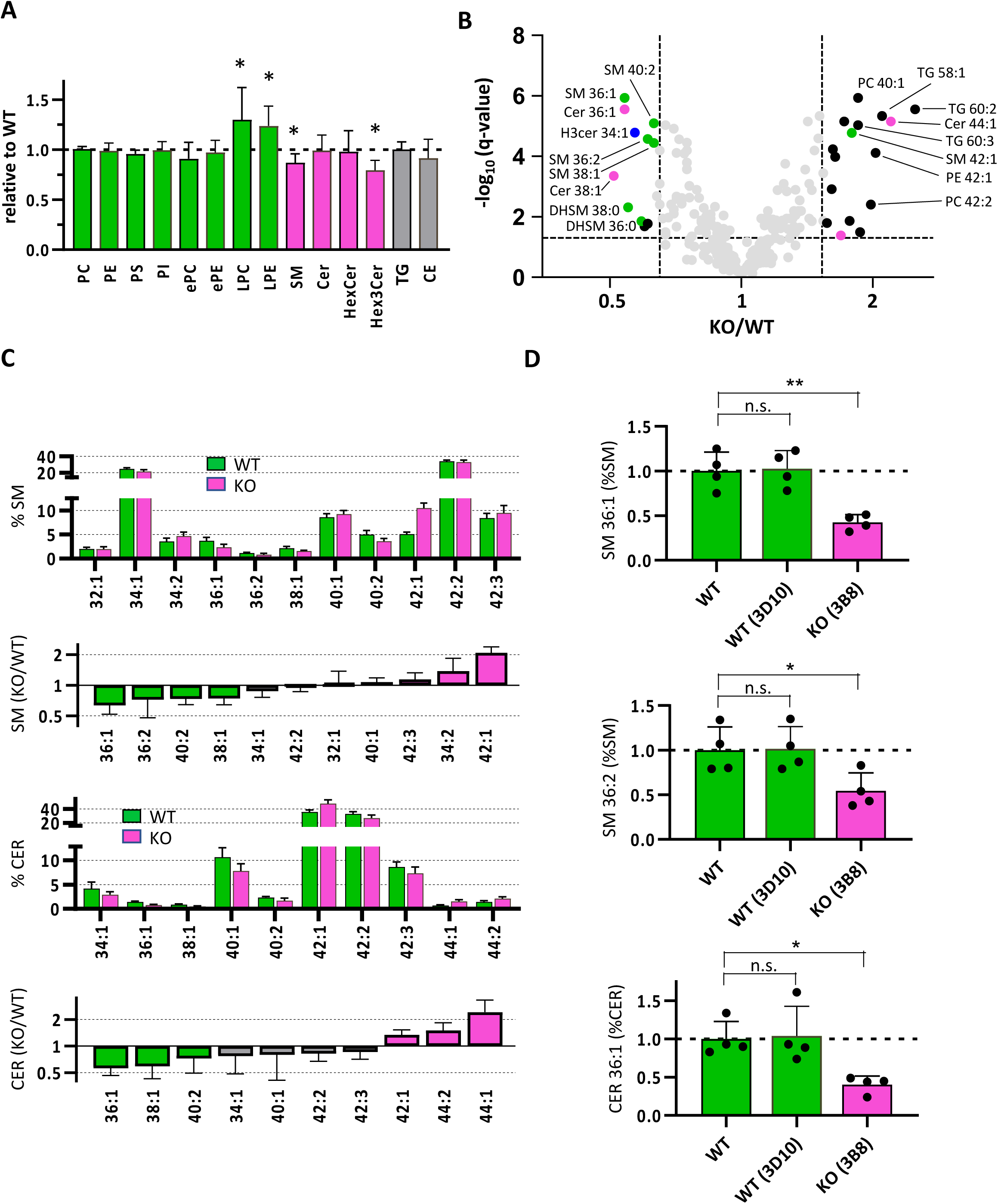
Reduction of sphingolipids with C18 acyl chains in VPS13B KO cells. **A)** Abundance of lipid classes in VPS13B KO cells (clone 2D9) relative to WT. Glycerophospholipids (green): phosphatidylcholine (PC); phosphatidylethanolamine (PE); Phosphatidylserine (PS); Phosphatidylinositol (PI); ether-PC (ePC); ether-PE (ePE); lyso-PC (LPC); lyso-PE (LPE). Sphingolipids (magenta): sphingomyelin (SM); ceramide (CER); hexosylceramides (HexCer); trihexosylceramides (Hex3Cer). Neutral lipids (grey): triacylglycerol (TG); cholesteryl ester (CE). (*p<0.01. paired t-test, 2-tails; N=12). **B)** Volcano plot illustrating relatives changes in individual lipid species in VPS13B KO cells (clone 2D9) relative to WT. sphingomyelins (green); ceramides (magenta); other lipid classes (black). (N=12). **C)** Relative abundance of individual SM and CER species expressed as a percentage of the respective class (upper charts) or as a ratio between VPS13B and WT amounts (N=12) **D)** Relative amounts of SM 36:1, SM 36:2 and CER 36:1 (calculated as a percentage of the respective class) in WT parental cells (WT), in WT cells (clone 3D10) and in VPS13B KO cells (clone 3B8). (*p<0.05, **p<0.01. Brown-Forsythe ANOVA with Dunnett T3 correction; N=4)

### Drug treatment partially corrects lipid defects in VPS13B KO cells

Since several CAD correct Golgi morphology defects in CS cells, we investigated whether these drugs could also revert lipid defects in VPS13B KO cells. To this end, we performed lipid analysis on cells treated with six different compounds: dilazep, azelastine, raloxifene, trifluoperazine, cyclobenzaprine and sertraline. As expected from the lysosomotrophic properties of these molecules, a 24-hour treatment led to a substantial accumulation of several lipid classes mostly sphingolipids, lysophospholipids and cholesteryl esters. Strikingly, different compounds exhibited highly heterogeneous lipid accumulation patterns, likely reflecting their selective inhibition of different lysosomal lipases. Sertraline, for instance, causes a pronounced accumulation of hexosylceramides (HexCer) without detectable storage of SM, whereas azelastine, trifluoperazine and cyclobenzaprine displayed an opposite preference. Additionally, sertraline, and to a lesser extent azelastine, trifluoperazine and cyclobenzaprine, also cause a significant increase in phosphatidylinositol (PI) levels. Dilazep, on the other hand, causes only limited storage of both SM and HexCer, but preferentially led to lysophospholipid (LPC and LPE) buildup (Fig. 7A, File S4). Other lipid classes are only marginally affected by compound treatment or even decreased as observed for PE, Hex3Cer and TG for a subset of compounds (Fig. S14). When analyzing individual lipid species, we observed that although most ceramide and sphingomyelin (SM) forms were more elevated in azelastine-and trifluoperazine-treated cells, certain species, including C18 acyl chain-containing forms (36:1, 36:2), which were depleted in VPS13B knockout (KO) cells compared to wild-type (WT) cells, showed significant enrichment following drug treatment (Fig. 7B, File S4). Furthermore, analysis of SM and ceramide distribution revealed that the same lipid species were enriched across all six treatment conditions (Fig. S15), with SM 36:1 reaching levels close to the WT condition after a 24-hours treatment (Fig. 7C). CER 36:1 instead reaches levels up to 4 folds higher than WT levels (Fig. S15B). Notably, species that were elevated in VPS13B KO cells (SM 42:1; CER 44:1) exhibited a relative reduction following drug treatment (Fig. S15), although their absolute levels remained almost unchanged (Fig. 7B, File S4). Additionally, the minor lipid species, which were elevated in VPS13B KO (PC 40:1; PE 42:1; TG 58:1, TG 60:2, TG 60:3), with few exceptions, are now reduced upon compound treatment (Fig. S16), which seems then to revert the VPS13B KO effect on lipid perturbations.

**Figure 7.**
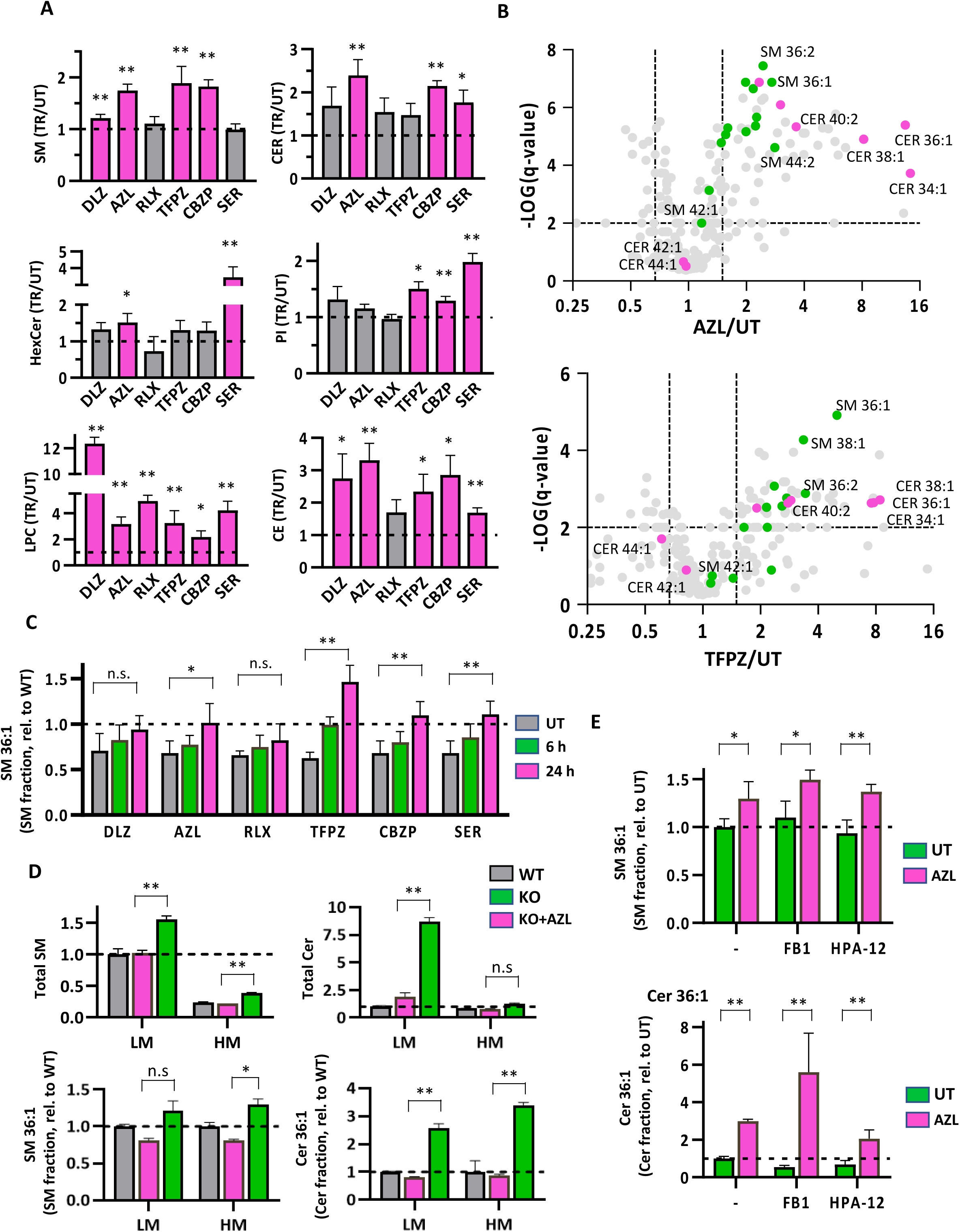
Drug treatment corrects lipid defects in VPS13B KO cells. **A)** Abundance of indicated lipid classes in VPS13B KO cells (clone 2D9) treated with the indicated compounds (10 µM, 24 hours), relative to untreated cells. DLZ (dilazep); AZL (azelastine); RLX (raloxifene); TFPZ (trifluoperazine); CBZP (cyclobenzaprine); SER (sertraline). (*q<0.05, **q<0.01. multiple paired t-tests with Benjamini-Hochberg correction; N=4) **B)** Volcano plot illustrating relatives changes in individual lipid species in compound treated VPS13B KO cells (clone 2D9) (**upper panel**, AZL; **lower panel**, TFPZ) relative to WT. Cells at 10 µM for 24 hours. sphingomyelin (green); ceramides (magenta) are highlighted. (AZL (N=12); TFPZ (N=6). **C)** Relative amounts of SM 36:1 (calculated as a fraction of total SM) in VPS13B KO cells treated with the indicated compounds for 6 or 24 hours (10 µM). (*q<0.05, **q<0.01. one-way ANOVA with Šidák correction; N=4) **D)** Relative amounts of total SM, total CER, SM 36:1 (calculated as a fraction of total SM) and CER 36:1 (calculated as a fraction of total CER) in light (LM) or Heavy (HM) membranes prepared from WT or VPS12B KO cells treated (AZL) with not (UT) with AZL 10 µM for 24 hours. (*p<0.05, **p<0.01. paired t-test, 2-tails; N=2). **E)** Relative amounts of SM 36:1 and CER 36:1 (calculated as a fraction of the respective lipid class) in cells treated (AZL) or not (UT) with 10µM AZL for 6 hours in combination with 20 µM fumonisin B1 (FB1) or 5 µM HPA-12. (*p<0.05, **p<0.01. paired t-test, 2-tails; N=3).

The increase in C18 SM induced by lysosomotropic compounds could either reflect its accumulation in endo-lysosomes alongside other stored lipids or indicate an increase in other cellular areas, including the Golgi complex, where it would be more physiologically relevant. To distinguish between these possibilities, we performed subcellular fractionation of microsomal intracellular membranes using a well-established protocol^47–49^. Although this method does not provide pure organelle fractions in HeLa cells, it effectively separates most endo-lysosomal membranes with a relatively minor TGN contamination in the fraction referred here as light membranes (LM). The cis-Golgi and ER are instead exclusively recovered in the heavy membrane (HM) fraction, which also contains the majority of the TGN (Fig. S17).

We prepared subcellular fractions from control and AZL-treated cells. As expected, SM and ceramides accumulated almost exclusively in the light membrane (LM) fraction following drug treatment. The same pattern was observed for other lipid classes prominently stored upon drug treatment, such as LPC and CE (Fig. 7D, upper panels; Fig S18). Similar results were obtained in TFPZ-treated cells (Fig.S19).

However, when analyzing the distribution of SM and CER species, we observed that C18 species (SM 36:1 and CER 36:1) also increased in the HM fraction, with SM 36:1 increasing to a similar extent as in LM, and CER 36:1 even in a more important manner (Fig. 7D, lower panels).

These results could suggest that the elevation of C18 sphingolipids may result from a metabolic regulation in response to lipid storage rather than merely from preferential accumulation in lysosomes. To test this hypothesis, we stimulated the cells with AZL for 6 hours in the presence of either the Ceramide Synthase (CERS) inhibitor fumonisin B1 (FB1)^50^ or the Ceramide transfer protein (CERT) inhibitor HPA-12^51^. If the observed increases in SM 36:1 and CER 36:1 were due to newly induced synthesis, FB1 would be expected to prevent both increases, while HPA-12 would selectively block the increase of SM 36:1. As shown in figure 7A, neither inhibitor was able to prevent C18 sphingolipid accumulation, excluding that the increase may indeed be due to enhanced synthesis. As expected, FB1 caused a significant reduction in total ceramide levels, while HPA-12 reduced total SM levels (Fig. S20).

The effect of CAD on Golgi recovery is phenocopied by genetic induction of lysososomal lipid storage, we wished to verify if also depletion of NPC1 or SMPD1 could also result in an increase of C18 sphingolipid levels. Given the availability of several lipidomic datasets modelling LSD in cell, we decided to re-analyse them in order to specifically focus on the variation in C18 sphingolipids. We analyzed five different datasets produced in either NPC1^52–54^-of SMPD1^54–56^-depleted cells either with KO or KD models. Four out of six were made in HeLa cells, the other two in HEK293 and A673 cells. Since LSD may cause a bulk accumulation of sphingolipids, similarly to our own work, we expressed the C18 species as a percentage of total class in order to highlight only specific variations. Despite heterogeneous experimental conditions in the different datasets, the relative levels of SM 36:1 are increased in all datasets for both NPC1 and SMPD1 depletion (Fig. S21A, B, File S5) with fold increases ranging from 1.5 to 3. CER 36:1, being a less abundant lipid is either not detected or very close to detection limit in most datasets. Nevertheless, when a measure is available ^53^, it also increases by 1.8 folds upon NPC1 KO (Fig. S21A). The study of Kraus et al. includes the proteo-lipidomic profiling of a more than LSD mutants allowing a broader correlation between lysosomal storage and C18 SM increase. Out of the 33 mutants, 17 cause a significant increase in SM 36:1 (File S5) including *Niemann-Pick C2* (NPC2) and proteins involved in sphingolipid degradation such as *Beta-hexosaminidase subunit alpha* (HEXA), *Prosaposin* (PSAP) and *Acid ceramidase* (ASAH1) (Fig. S21C).

### Azelastine and raloxifene partially restore neurite outgrowth in primary neuronal cultures derived from cortical organoids

We recently developed a novel culture protocol to generate brain organoids from human pluripotent stem cells (hPSCs). These organoids recapitulate key aspects of anterior central nervous system (CNS) development and exhibit cellular diversity and transcriptomic profiles that closely resemble those of the developing human cerebral cortex^57^. The neural organoids derived from VPS13B KO hPSCs, grow at normal growth rate compared to control organoids until 1 month of age. However, at 2-month stage VPS13B KO organoids are significantly smaller than WT controls^57^. This reduction in size is not due to decreased cell proliferation or increased apoptosis, but rather to smaller soma size and impaired neurite outgrowth—findings consistent with previous reports in primary hippocampal neurons^14^. Collectively, these findings support the utility of VPS13B KO brain organoids as a faithful in vitro model of the secondary microcephaly observed in Cohen syndrome (CS) patients.

To evaluate the potential of CAD in rescuing developmental defects, we tested two candidate compounds, azelastine and raloxifene, for their ability to restore the size of organoids as well as the neurite outgrowth in VPS13B KO neurons. We initially tested both compounds at a 10 µM concentration noticing an evident drug-induced toxicity on both WT and VPS13B KO organoids over the long-term treatment required for these experiments (30 days). We performed a toxicity titration on WT organoids, fixing the final working concentration at 1 µM on in intact organoids and at 500 nM in isolated neurons. Primary neurons were prepared from one-month-old organoids and cultured for three days in the presence or absence of either compound. Remarkably, both azelastine and raloxifene significantly promoted neurite outgrowth in VPS13B-deficient neurons (Fig. 8A–D). To assess whether these compounds could also rescue the reduced size phenotype of intact organoids, treatments were administered from day 25 to two months of age. In this context, however, neither compound was able produce a significant effect in recovering organoid size (Fig. 8E–H).

**Figure 8.**
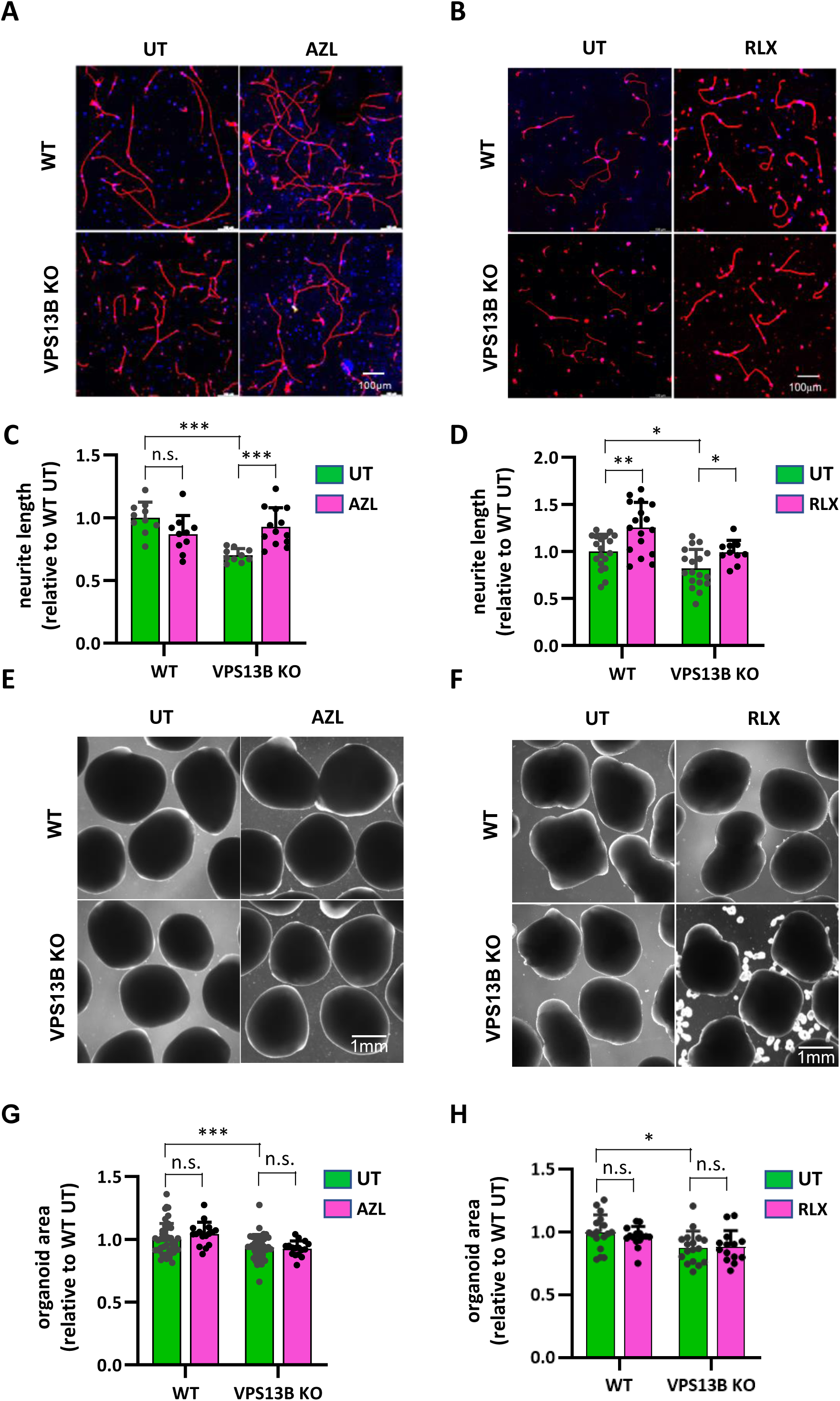
Azelastine and raloxifene restore neurite outgrowth in neurons. A, **B)** Representative images of control and VPS13B KO neurons treated with 500 nM azelastine (AZL) and raloxifene (RXL). Cells stained with Hoechst (blue) and Tuj1 (red). Scale bar: 100 μm. **C, D)** Quantification of mean neurite length in control and VPS13B KO neurons following AZL or RLX treatment. Neurite length values were normalized to DMSO-treated control neurons. Each data point represents the mean neurite length of approximately 9–16 individual neurons. Data are presented as mean ± SD. (Welch ANOVA test with Dunnett T3 multiple comparisons correction; ***p<0.001; **p<0.01; *p<0.05; ns: non-significant). **E, F**) Representative brightfield images of control (top) and VPS13B KO neural organoids (bottom) at the 2-month stage, treated with 1 µM of AZL and RLX for 35 days. Scale bar: 1 mm. **G, H**) Quantification of organoid area following AZL and RLX treatment. Organoid sizes were normalized to untreated control organoids. (Welch ANOVA test with Dunnett T3 multiple comparisons correction; ***p<0.001; **p<0.01; *p<0.05; ns: non-significant; n= ∼16 organoids).

## DISCUSSION

In this study, we screened a well-annotated library of 1280 compounds, primarily FDA-or EMA-approved drugs and identified several molecules capable of efficiently restoring Golgi morphology in VPS13B KO cells. A detailed analysis of target specificity, chemical properties, as well as dose-response results, led us to the conclusion that for the vast majority of the compounds the observed biological effect was unlikely to result from direct interaction with a specific molecular target. Instead, the mechanism appeared to involve lipid accumulation in acidic organelles induced by compounds with CAD properties, a phenomenon known as drug-induced phospholipidosis^24^. This conclusion was further supported by the observation that genetically induced lipid storage, achieved by NPC1 or SMPD1 silencing, similarly restores Golgi complex morphology.

The CADs were unable to prevent Golgi disassembly caused by either BFA-mediated ARF1 inhibition or microtubule depolymerization. Additionally, they did not accelerate Golgi recovery after the washout of BFA or nocodazole. These observations suggest a degree of specificity in the effect of the compounds and highlight the dependence of their action on microtubule integrity. To the contrary, the dramatic Golgi dispersion induced by statins, as identified in our screening and consistent with previous findings^39^, was completely reversed by CADs. A more comprehensive interpretation of these outcomes will likely emerge once the mechanistic basis of Golgi structure perturbation in CS is elucidated, which may also enable the assessment of potential similarities with other perturbation triggers.

Our findings also indicate that the more remarkable lipid alteration in VPS13B KO cells is the decrease in both SM and ceramides with 18 and 20 carbons acyl chains, observed both in absolute values and relative to the corresponding lipid class. C18 species in particular, were originally shown to be highly enriched COPI vesicles^42^ and later found to specifically bind and regulate TMED proteins^43^; ^44^. Interestingly, loss of TMED2/10 has also been associated with Golgi complex dispersion^58^. Taken together these observations provide an intriguing link with the Golgi phenotype of VPS13B-deficient cells, although a direct cause-effect relationship between Golgi fragmentation and reduced C18 SM levels cannot yet be firmly established.

The treatment with six different CADs, selected from the hits of our screening, leads to a significant increase in C18-20 sphingolipid species, even when normalized to their respective lipid classes. Notably, while C18 ceramide levels rise well above those observed in WT cells, C18 SM levels return close to WT values following drug treatment. Interestingly, cell fractionation experiments reveal that C18 SM and ceramide levels increase significantly also in the ER/cis-Golgi HM fraction, whereas bulk lipid accumulation is restricted to the endo/lysosome-rich LM fraction. Moreover, results with the CERS and CERT inhibitors indicate that the rise in C18 species is unlikely to result from upregulated synthesis of these lipids in response to lysosomal storage, since their increase was not prevented in the presence of these inhibitors. The increase in C18 sphingolipids in response to CAD is not entirely novel since it was already observed in the hippocampus of mice treated with CAD antidepressants. In this study, C18 sphingolipids are also shown to increase in the ER-rich fraction and not exclusively in lysosomes^59^. Intriguingly, re-analysis of published lipidomic datasets from NPC1-and SMPD1-depleted cells also reveals an increase in C18 sphingolipids, reinforcing the notion that genetic silencing of these genes and CAD treatment may operate through a shared mechanism of action. Notably, this analysis further indicates that lysosomal storage–induced increases in these sphingolipid species also occur in WT cells, suggesting that lysosomal storage does not restore VPS13B function in C18 sphingolipid homeostasis per se. Rather, lysosomal lipid storage appears to generally elevate the relative levels of these species, potentially by reducing their lysosomal turnover. Such a mechanism could indirectly compensate for their depletion in CS cells.

The concept of lysosomotropic drugs for CAD molecules was first introduced by Christian de Duve half a century ago^60^. Subsequent studies revealed that these compounds accumulate in acidic organelles, where they inhibit lipid hydrolase activity by disrupting interactions with the negatively charged intraluminal membranes^24^; ^61^; ^62^. CAD molecules have been shown to inhibit multiple enzymes, including acidic sphingomyelinase^62^, ceramidase^63^, and phospholipases A1^64^ and A2^65^. The heterogeneous lipid storage patterns observed in our study likely reflect the selective inhibition of specific enzymes by different compounds.

In current drug discovery programs, lysosomotropism is often regarded as a challenge in drug development because of its potential adverse effects. Nevertheless, several CAD have been successfully approved and remain in clinical use. More recently, their potential repurposing for other diseases has been also proposed, leveraging their ability to modulate lipid metabolism, particularly sphingolipid pathways.

Studies in mouse models have shown that tricyclic antidepressants may exert their biological effects, at least in part, by modulating sphingomyelin and ceramide levels in the brain^59^; ^66^. Additionally, several CADs have been identified in a screening as potential inhibitors of hepatic fibrosis ^67^ and amitriptyline, a tricyclic CAD also identified in our study, has been shown to mitigate high-fat diet-induced steatohepatitis in mice ^68^. In both cases, the therapeutic effects appear to be mediated by sphingolipid regulation.

More recently, Soldati et al. proposed repurposing tamoxifen for the treatment of two forms of ceroid lipofuscinosis through a mechanism involving lysosome-dependent activation of the TFEB transcription factor^69^. CADs are also under active investigation in cancer therapy, particularly for their ability to modulate lysosomal lipid levels, inducing tumor cell-selective toxicity^70–72^.

In this study we found that several CAD restore Golgi morphology in HeLa cells and two compounds were able to partially rescue neurite outgrowth in primary neurons dissociated from a brain organoid CS model that recapitulates the postnatal microcephaly phenotype observed in patients. This finding is particularly significant, as it indicates that the therapeutic effects of these compounds extend beyond Golgi structural correction to include functional neuronal recovery in a physiologically relevant disease model. Notably, the long-term treatment regimen required lower drug concentrations than those used for acute experiments in HeLa cells. This observation underscores the distinctive mechanism of action of these compounds, which progressively accumulate within intracellular organelles over time, and raises important considerations regarding their dosing and optimal use *in vivo*.

While we recognize the need for caution regarding the therapeutic use of lysosomotropic compounds, our findings, consistent with studies cited above, highlight the potential to identify lipid regulatory pathways, representing novel potential therapeutic targets for CS. Gaining further mechanistic insights will be crucial to uncover the precise mechanism of action of CADs in this context and design more selective drugs that could correct the disease phenotype without the potential drawback of-massive lysosomal lipid storage. Moreover, we believe that our results also reinforce the value of phenotypic compound screenings, not only as a means to identify therapeutic protein targets, but also to uncover novel potential intervention mechanisms.

## MATERIALS AND METHODS

### Cell culture and transfections

HeLa-MZ cells were authenticated by Microsynth (Balgach, Switzerland), showing 100% identity with the DNA profile of HeLa (ATCC CCL-2) and full concordance across all 15 autosomal short tandem repeats (STRs) with Microsynth’s HeLa reference profile. Cells were maintained in Dulbecco’s Modified Eagle Medium (DMEM) with high-glucose (4.5 g/L) supplemented with GlutaMAX, 10% fetal bovine serum (FBS), and penicillin-streptomycin. Transfection with siRNA was performed as previously described^26^. Briefly, cells were seeded in 6-well plates 6 h before transfection in antibiotic-free medium (3×10^5^ cells/dish). Cells were then transfected with siRNA max (5.25 µl/well; Thermo Fisher Scientific) and siRNA (70 pmol/well) following the manufacturer’s instructions. After 48 hours, cells were harvested and plated into 96-well imaging plates (Ibidi, Gräfelfing, Germany; ref. no. 89626) at a density of 1.2×10⁴ cells per well. Fixation and imaging were carried out 24 hours later (72 hours post-transfection). The following siRNAs were used are: NPC1 (Qiagen, cat. no. Hs_NPC1_4) and SMPD1 (siTOOLs BIOTECH, siPOOL (gene ID: 6609). For Halo-VPS13B plasmid (generous gift of Pietro De Camilli) transfection, cells were seeded at a density of 8×10^3^ cells per well into 96-well imaging plates in antibiotic-free medium. After 15 hours, cells were transfected with Fugene 4K (0.9 µl/well; Promega) and Plasmid DNA (0.3 µg/well). The culture medium was replaced 5 hours after transfection and cells were fixed 32 hours after transfection.

### Antibodies, Dyes and Chemicals

Primary antibodies used in this study are: Calnexin mouse mAb clone 6F12BE10 (Abcam, cat. ab112995); Early endosomal antigen 1 (EEA1) rabbit pAb (Enzo Life Sciences, cat. ALX-210-239); GM130 mouse mAb clone 35 (BD Transduction, cat. 610823); GOLPH3 rabbit pAb (Abcam, cat. ab98023); HaloTag rabbit pAb (Promega, cat. G9281); LAMP1 rabbit mAb clone D2D11(Cell Signaling, cat. 9091); LBPA mouse mAb clone 6C4^73^ (generous gift of Jean Gruenberg); Perilipin 2 (PLIN2) mouse mAb clone AP125 (Progen, cat. 610102); Transferrin Receptor mouse mAb clone H68.4 (Thermo Fisher, cat. 13-6800); VPS13B rabbit pAb (Atlas antibodies, cat, HPA043865).

Secondary antibodies used in this study are: Donkey Anti-Mouse IgG (H+L) (Jackson Immunoresearch, cat. 715-545-150 (Alexa Fluor 488); 715-585-150 (Alexa Fluor 594); 715-605-150 (Alexa Fluor 647)); Donkey Anti-Rabbit IgG (H+L) (Jackson Immunoresearch, cat. 711-545-152 (Alexa Fluor 488)).

Fluorescent dyes used in this study are: Hoechst 33342 (Thermo Fisher, cat. 62249); Propidium iodide (Thermo Fisher, cat. P1304MP); HCS LipidTOX™ Green Phospholipidosis Detection Reagent (Thermo Fisher, cat. H34350); Filipin III (Merck, cat. F4767).

Prestwick chemical library compounds used in the primary screening and dose-response curves were purchased from Prestwick chemical libraries (Orléans, France). Catalog ID and compound details are indicated in File S1. Compounds used in further follow-up experiments were purchased separately: azelastine hydrochloride (MedChemExpress, cat. HY-B0462 for HeLa cells experiments; Sigma-Aldrich, cat. A7611 for brain organoid experiments); brefeldin A (Merck, cat. B6542) cyclobenzaprine hydrochloride (Merck, cat. T8516); dilazep dihydrochloride (MedChemExpress, cat. HY-100957); fumonisin B1 ((Merck, cat. F1147); HPA-12 (MedChemExpress, cat. HY-132182); lovastatin (MedChemExpress, cat. HY-N0504); nocodazole (Apex Bio, cat. A8487); raloxifene (MedChemExpress, cat. HY-13738 for HeLa cells experiments; Sigma-Aldrich, cat. PHR1852 for brain organoid experiments); sertraline hydrochloride (Merck, cat. S6319); trifluoperazine dihydrochloride (Merck, cat. C4542); U18666A (Merck, cat. U3633).

### CRISPR KO cell line generation

VPS13B knockout (KO) HeLa cells were generated via CRISPR/Cas9-mediated genome editing by introducing small insertions or deletions (INDELs) in the open reading frame (ORF) of the VPS13B gene, as previously described^74^ with minor modifications. Target genomic sites within exons 2 and 3 of VPS13B gene (File S6) were selected using CRISPOR online tool^75^ (https://crispor.gi.ucsc.edu/). Sites were prioritized based on high on-target cleavage efficiency and low off-target potential^76^, as well as high out-of-frame mutation likelihood using the LINDEL score^77^. Oligo DNA pairs were synthesized, annealed and cloned in px330 vector^78^ (Addgene #42230) as described^74^. HeLa MZ cells were plated in 6 cm dishes (7.5 x 10^5^ cells/plate) and transfected after 18 hours using Lipofectamine 3000 (6 μL/dish; Thermo Fisher Scientific) with 4 μg plasmid DNA per dish. Five days post-transfection, single cells were isolated using a SH800S Cell Sorter (Sony Biotechnology) and plated into 96-well plates (one cell per well; three plates per condition) to establish clonal populations. After clone expansion, genomic DNA was extracted with Wizard Genomic DNA Purification Kit (Promega corporation) and the target genomic regions were PCR-amplified. Amplicons were sequenced by Sanger sequencing (Fasteris SA, Plan-les-Ouates, Geneva, Switzerland) with primers internal to the amplicon sequence (File S6 for primer sequences). INDELs resulting in frameshift mutations were confirmed via chromatogram decomposition using the TIDE online tool^79^ (https://tide.nki.nl/) (Fig. S1A).

### SDS-PAGE electrophoresis and Immunoblotting

Protein concentration was determined using the Bradford assay. For comparison of sub-cellular fraction, equal volumes of each fraction were diluted in 4× Laemmli loading buffer and resolved by SDS-PAGE. 10 μg of protein from the heavy membrane (HM) fraction were loaded per lane, corresponding to 1.5-2 μg for LM fraction. 25 μg of proteins from post-nuclear supernatant (PNS) were loaded as a control. Following electrophoresis, proteins were transferred to membranes (Amersham Hybond P PVDF membrane) and probed with primary antibodies (VPS13B rabbit pAb 1:100; LAMP1 rabbit mAb 1:1000; EEA1 rabbit pAb 1:200; GOLPH3 rabbit pAb 1:1000; GM130 mouse mAb 1:500; Calnexin mAb 1:500) overnight at 4 °C. HRP-conjugated secondary antibodies were incubated for 50 minutes at room temperature. Detection was performed using WesternBright Quantum chemiluminescent substrate (Advansta, Menlo Park, CA) and visualized with the FUSION Solo imaging system.

### High-content microscopy Screening

The compound library was pre-spotted into 384-well imaging plates (Ibidi, Gräfelfing, Germany; cat. no. 88416) at a final amount of 0.5 nmol/well (50 nL of 10 mM stock in DMSO) by the Biomolecular Screening Facility (EPFL, Lausanne, Switzerland) in order to get a concentration of 10 µM after cell seeding. Each plate included four columns of vehicle-only control wells. Wild-type (2 control columns per plate) and VPS13B knockout HeLa-MZ cells (2 control columns per plate plus compound wells) were seeded at a density of 2,000 cells per well in 50 µL of complete medium using a Multidrop Combi+ liquid dispenser (ThermoFisher Scientific). After 24 hours, cells were fixed with 50 µL of 3% (w/v) paraformaldehyde (PFA) in PBS for 15 minutes at room temperature, then washed with PBS. For immunostaining, cells were incubated with a primary antibody (GM130; 1:500) in 20 µL PBS with 0.05% saponin and 1% bovine serum albumin (BSA). After washing, cells were incubated with Alexa Fluor 488-conjugated anti-mouse secondary antibody (1:200) and Hoechst 33342 (1:2000) in 20 µL PBS. Antibody dispensing and washing steps were performed using a Biotek EL406 plate washer. Images were captured using an ImageXpress Micro Confocal High-Content microscope (Molecular Devices) equipped with a 40× objective. For each well, 25 fields were imaged across three z-stack planes. For DRC, experiments were conducted exactly as above with the exception of HCS LipidTOX Green phospholipidosis detection reagent (Thermo Fisher Scientific, cat. H34350) diluted in the plating medium (1:2000) and the use of Alexa Fluor 647-conjugated anti-mouse (1:200) as secondary antibody to detect GM130.

### Other immunofluorescence experiments

For all other experiments with single or double antigen detection cells were fixed in 96-well imaging plates with 70 µL of 3% (w/v) PFA in PBS. For immunostaining, cells were incubated with a primary antibodies (GM130 mouse mAb 1:500; GOLPH3 rabbit pAb 1:500; HaloTag rabbit pAb 1:1000; LAMP1 rabbit mAb 1:1000; TFR mouse mAb 1:300; PLIN2 mouse mAB 1:100; LBPA mouse mAb 1:100) in 70 µL PBS with 0.05% saponin and 1% bovine serum albumin (BSA) for 1 hour. After washing, cells were incubated appropriate secondary antibodies (Alexa Fluor 488-anti-rabbit; Alexa Fluor 488-anti-mouse; Alexa Fluor 594-anti-mouse; Alexa Fluor 647-anti-rabbit;1:200) with Hoechst 33342 (1:2000) in 70 µL PBS. All washing steps were performed using a Biotek EL406 plate washer. Images were captured using an ImageXpress Micro Confocal High-Content microscope (Molecular Devices) equipped with a 20X or 40X water immersion objectives. For each well, 6-9 fields (20X objective) or 25 fields (40X objective) were imaged across three z-stack planes. Filipin staining was performed as previously described^26^, pre-treating fixed cells with RNase (200 μg/ml) followed by propidium iodide (PI) (5 μg/ml) for nuclei staining and filipin (50 μg/ml) in PBS.

### Image analysis and quantification

Image analysis was conducted using the MetaXpress Custom Module editor software (Molecular Devices). Cell segmentation was performed as previously described^80^ based on Hoechst 33342 signal intensity. Two different intensity thresholds were applied to distinguish between nuclear and cytoplasmic regions. Cells undergoing mitosis were identified by their high Hoechst 33342 intensity in condensed chromosomes and were excluded from analysis. Cells located at the image borders were also excluded from analysis.

Golgi morphology was quantified by binarizing the GM130 signal using a defined threshold to generate a Golgi mask, enabling extraction of the following features: average area and perimeter of Golgi fragments, number of fragments per cell, and shape factor. To enable compound ranking, parameter values were first standardized. We then calculated the 4-dimensional Euclidean distance between average well values of untreated wild-type (WT) and knockout (KO) controls (n = 32 wells per plate), across the four parameters:

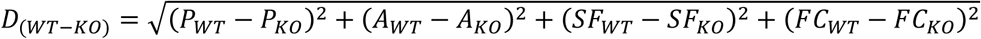

Where: (P=perimeter; A=area; SF=shape factor; FC=fragments/cell)

This WT–KO distance was normalized to 1.0. For each test well, the distance to WT was calculated using the same formula. The interpretation of distance values (D) with respect to Golgi morphology recovery is then as follows: (D=0 full recovery; 0<D<1 partial recovery; D=1 no effect; D>1 deleterious effect).

All calculations were performed using plate-specific internal controls. Same threshold settings were used within each experimental batch. Wells with a cell count below 67% of the plate average were considered potentially cytotoxic and excluded from the final ranking.

For experiments involving transfection with the Halo-VPS13B construct (Fig. 1D), fluorescence intensity in the Halo channel was quantified alongside Golgi morphology parameters detected via the GM130 channel. A defined intensity threshold for the Halo signal (established using mock-transfected control cells) was applied to exclude non-transfected cells from analysis. Additionally, cells exhibiting excessively high Halo-VPS13B expression were excluded to minimize potential artifacts from overexpression.

### Sub-cellular fractionation

Subcellular fractions were prepared by sucrose gradient flotation as previously described^47–49^ with minor modifications. Briefly, cells grown in 15 cm dishes (3 dishes per condition) were harvested by gentle scraping in ice-cold PBS using a flexible rubber policeman to obtain sheets of attached cells to minimize mechanical disruption. Cells were pelleted at 175 g for 5 min at 4°C, resuspended in homogenization buffer (HB) (250 mM sucrose, 3 mM imidazole, pH 7.4), and pelleted again at 1,300 g for 7 min. Cell pellet was resuspended in 0.5–1 ml HB and homogenized by passage through a 22-gauge needle using a 1-ml syringe. The homogenate was centrifuged at 1,300 g for 7 min. The supernatant was collected, adjusted to 40.6% sucrose, loaded at the bottom of an ultracentrifugation tube, and overlaid sequentially with 35% sucrose in 3 mM imidazole (pH 7.4), and then with HB. Gradients were centrifuged at 165,000 g for 1.5 h at 4°C with a SW40Ti Swinging-Bucket rotor (Beckman-Coulter). The LM and HM fractions were collected at the 35%-HB and 35%-40.6% interfaces, respectively.

### Human PSCs culture and neural organoid generation and treatment

H9 human pluripotent stem cells (hPSCs) were cultured on Matrigel-coated plates (Corning, NY, USA) in mTeSR1 medium (STEMCELL Technologies, Vancouver, Canada). Cells were passaged every 4–5 days using ReLeSR (STEMCELL Technologies) following the manufacturer’s protocol.

Neural organoids (NOs) were generated as described^57^. Both control and VPS13B KO H9 hPSCs were used. Neural induction was initiated via dual SMAD inhibition with SB431542 (10 uM, Tocris, Cat. #1614) and LDN193189 (100 nM, Stemgent, Cat. # 04-0074) in differentiation medium (DM) for 3 days with daily medium changes. The DM is composed of DMEM/F12 (Gibco, Cat. #11320-33), 2% B-27 Supplement (Gibco, Cat. #17504044), 1% N-2 Supplement (Gibco, Cat. #17502-048), 1% MEM Non-Essential Amino Acids (Gibco, Cat. #11140-050), 1% penicillin-streptomycin (Gibco, Cat. #15140148), and 0.01% β-mercaptoethanol (Gibco, Cat. #21985-023). Following induction, neural stem cells were dissociated into single cells using accutase (STEMCELL Technologies) and seeded at a density of 5,000 cells per well in a 96-well ultra-low attachment plate. Cultures were maintained in DM supplemented with 20 ng/mL basic fibroblast growth factor (bFGF) and 20 ng/mL epidermal growth factor (EGF) for 4 days, with daily supplementation. Subsequently, aggregates were cultured in DM without growth factors for an additional 8 days, with medium changes every other day.For maturation, NOs were transferred to a 1:1 mixture of DMEM/F12 and Neurobasal medium, supplemented with 2% B-27, 0.5% N-2, 0.5% MEM NEAA, 1% penicillin-streptomycin, 1% GlutaMAX (Life Technologies), and 0.1% β-mercaptoethanol. The medium was refreshed every three days. For compound treatment, azelastine and raloxifene were dissolved in DMSO at 10 mM stock concentrations and diluted in culture medium to the desired final concentrations. Organoids were treated with drug-containing medium every three days from day 25 to the two-month stage. Untreated organoids receiving DMSO only served as vehicle controls. Brightfield images were captured every three days using an EVOS M5000 Imaging System, and organoid size was quantified using ImageJ software.

### Primary neuron culture from organoids and treatment

Primary neurons were derived from 1-month-old organoids by enzymatic dissociation using the Papain Dissociation System (Worthington, Cat. #LS003126). Cells were plated onto Poly-L-Ornithine/laminin-coated coverslips at a density of 100,000 cells per well in a 24-well plate. The culture medium consisted of Neurobasal medium supplemented with 1% GlutaMAX, 2% B27 Supplement (Thermo Fisher), and 1% penicillin/streptomycin (P/S), 200 µM ascorbic acid phosphate (Sigma, Cat. #A8960), 125 µM cAMP (Sigma, Cat. #D0627), 1 µg/mL laminin (Invitrogen), 1% Culture One Supplement (Gibco, Cat. #A3320201), and 1% Chemically Defined Lipid Concentrate (Gibco, Cat. #11905031). Drug treatments were applied one day after plating and incubated for three days. DMSO was used as the vehicle control. After treatment, neurons were fixed with 4% PFA and stained for the neuronal marker Tuj1. Images were processed in ImageJ, and neurite length was manually measured using NeuronJ, (a plug-in for ImageJ). Total neurite length was calculated by summing the traced neurites from all cells within each image. Cell counts were obtained from the Hoechst channel, and the mean neurite length was determined by dividing the total neurite length by the number of cells. Only neurites within the field of view were traced, and non-neuronal cells (identified as Tuj1-negative nuclei) were excluded from the analysis. Drug-treated neurons were normalized to DMSO-treated control neurons

### Lipid extraction

Lipid extraction was carried out using a modified version of methyl tert-butyl ether (MTBE) protocol^81^, as previously described ^55^; ^82^ with minor modifications. Briefly, cells were seeded in 6-well plates (3.3×10^5^ cells/well) in complete medium containing 10% FCS and harvested after 24 hours for lipid extraction. Cells were first washed with cold PBS and scraped off with 460 μl cold methanol/H2O (3.6:1) on ice. The cell suspension was transferred to a 2.0 mL Eppendorf Safe-Lock Tube and a mixture of internal standards (File S7) was added. Subsequently, 1.2 mL of MTBE was added, and the samples were vortexed for 10 minutes at 4 °C, followed by a 1-hour incubation at room temperature with shaking at 750 rpm. Phase separation was induced by the addition of 200 μL of LC-MS-grade water and a 10-minute incubation. Samples were centrifuged at 1,000 × g for 10 minutes, and the upper organic phase was carefully transferred to a 2.0 mL amber glass vial. The remaining lower aqueous phase was re-extracted with 400 μL of artificial upper phase (MTBE/methanol/water, 10:3:1.5, v/v/v). The two organic phases were pooled and dried using a CentriVap concentrator at 55 °C. Finally, samples were flushed with nitrogen and stored at −80 °C until further analysis.

### Lipidomics Analysis by LC-MS

Dried samples were resuspended by sonicating in 100 µl of LC-MS-grade chloroform:methanol (1:1, v/v). Reversed-phase ultra-high-performance liquid chromatography–high-resolution mass spectrometry (UHPLC-HRMS) analyses were performed using a Q Exactive Plus Hybrid Quadrupole-Orbitrap mass spectrometer coupled to an UltiMate 3000 UHPLC system (Thermo Fisher Scientific) equipped with an Accucore C30 column (150 x 2.1 mm, 2.6 μm) and its 20mm guard column (Thermo Fisher Scientific). Samples were kept at 8°C in the autosampler, and 10 μl were injected for analysis. Lipids were separated using the following gradient: 10% solvent B for 1 min, ramped from 10% to 70% B over 4 min, then from 70% to 100% B over the next 10 min, held at 100% B for 5 min, followed by a 3-min re-equilibration. The mobile phases consisted of 5 mM ammonium acetate and 0.1% formic acid in water (solvent A) and in isopropanol:acetonitrile (2:1, v/v) (solvent B). The flow rate was maintained at 350 μL/min, and the column oven was set to 40 °C. The mass spectrometer was operated using a heated electrospray-ionization (HESI) source in positive and negative polarity with the following settings: electrospray voltage:-3.4 KV (-) or 3.9 KV (+); sheath gas: 51; auxiliary gas: 13; sweep gas: 3; vaporizer temperature: 431°C; ion transfer capillary temperature: 320 °C; S-lens: 50; resolution: 140,000; m/z range: 200-1500; automatic gain control: 1e6; maximum injection time: 50 ms.

### Data Processing and Lipid Quantification

Quantification of lipids was performed using Skyline v. 24.1.0.414 (MacCoss Lab, University of Washington, Seattle, USA)^83^; ^84^. For Skyline quantification, Orbitrap Raw files were converted into two separate mzML files for positive and negative ionization modes with ProteoWizard MSConvert software^85^. Transition lists of lipids were generated by using the LipidCreator Skyline-integrated tool^86^ and validated based of experimental data. Original transition list contains lipid class (group name), precursor name, precursor m/z, precursor formula and adduct. After technical validation, lipid species were assigned with explicit retention time (RT) values that were included in transition lists used for quantification with an indicated retention time window of 0.5 min. for individual lipid species (File S8). Only lipid species showing consistent RT behavior—according to established relationships between RT, acyl chain length, and degree of unsaturation in reversed-phase chromatography^87^, were retained for quantification (Fig. S22). Retention time alignment across LC-MS runs was performed using a reference set of 34 lipid species, with adjustments made directly in the Skyline transition list prior to quantification. All lipid classes were quantified in positive ion mode with the exception of PI and PS which were analyzed in negative ion mode. Peaks were extracted in skyline with a centroid m/z error tolerance of 2 ppm. Lipid species were considered valid for quantification only if their isotope dot product (idotp) exceeded 0.9. Quantification was based on peak height, which showed slightly better reproducibility than peak area across technical replicates. Peak height values were normalized to the internal standard of the corresponding lipid class Each lipid species is expressed as a normalized value to the total GP content in the sample. For lipid classes missing appropriate standard from our standard mix (CE, LPC, LPC), the values are reported without internal standard normalization. Due to the limited number of internal standards used, precise absolute quantification of lipid species and classes was not feasible. Therefore, all data are expressed as relative changes across experimental conditions rather than as absolute concentrations.

### Analysis of published lipidomic datasets

The published lipidomic datasets were analyzed starting from excel files published by the authors and relying on lipid quantifications performed by the authors. The analysis specifically focused on the relative quantification of SM and CER species expressed as a percentage of the corresponding class content. The studies focus on different types of biological questions and include several different treatment conditions. For the purpose of our study we only included untreated WT cell and LSD gene depleted cells. One study also includes sub-cellular fractionation experiments^52^. For consistency with the other studies, the lipid analysis obtained from total cells. The dataset from Kraus et al.^54^ is organized in different batches, each one with its own controls. For each gene KO, we used the controls within the same batch.

## ACKNOWLEDGEMENTS

The authors warmly thank Jean Gruenberg for his thoughtful reading of the manuscript and helpful comments. We are also grateful to Gerardo Turcatti and Marc Chambon for their valuable advice and technical input regarding compound screening. This research was supported by the Fondation Pro Visu (Geneva, Switzerland), which also provided salary support for FV. We gratefully acknowledge additional funding from the Swiss National Science Foundation (grants 320030_212959 and IZSTZ0_216615), AriBio Co., the Million Dollar Bike Ride grant program of the Penn Medicine Orphan Disease Center, the Prix Claire et Selma Kattenburg and Kun-hee Lee Seoul National University Hospital Child Cancer & Rare Disease Project, Republic of Korea (25B-01-0500).

---

**Figure S1.**
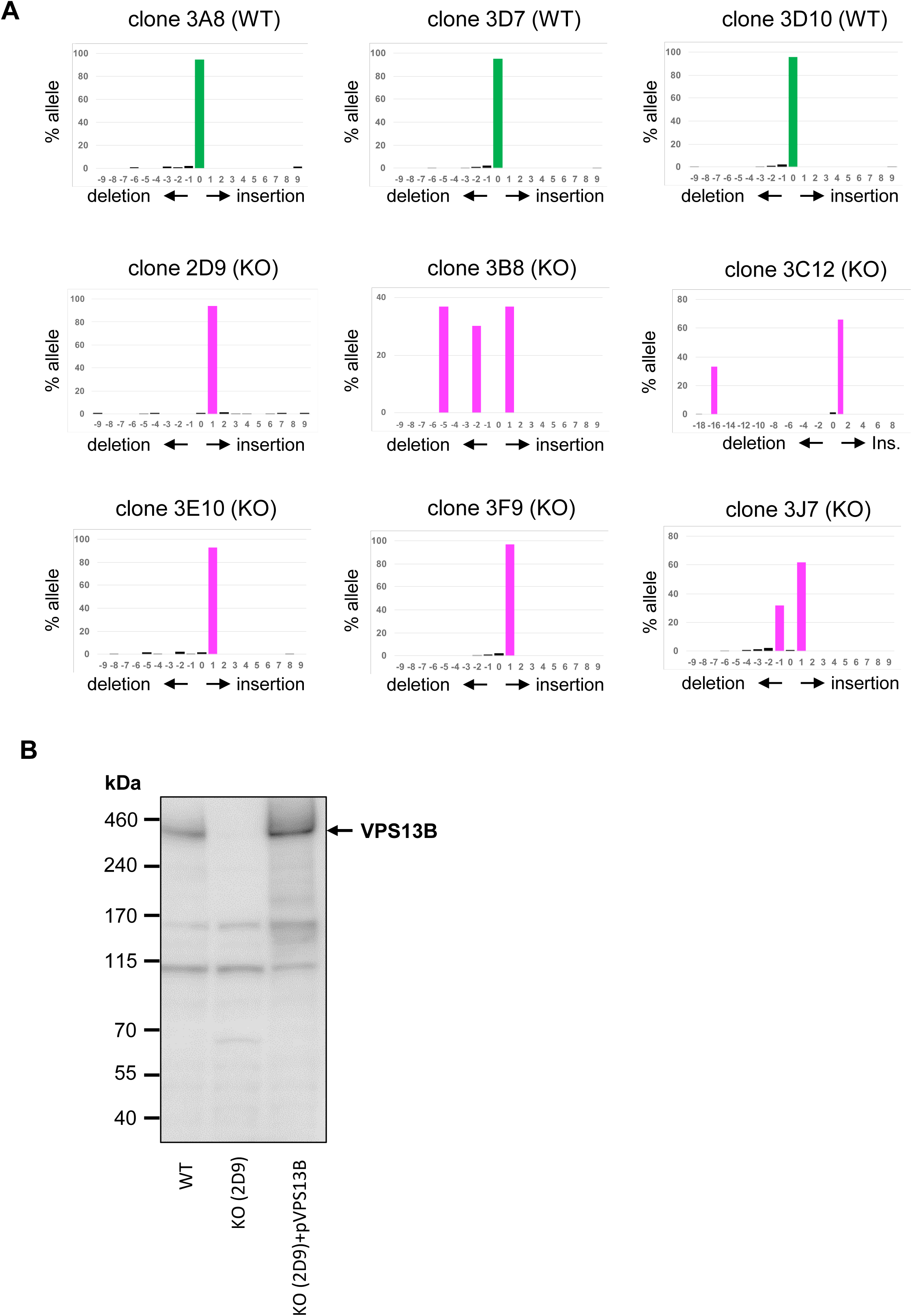
Validation of VPS13B knockout in HeLa cells. **A)** TIDE analysis of HeLa cell clones with either wild-type (WT) or homozygous VPS13B knockout (KO) genotypes, showing the distribution of insertions and deletions (INDELs) at the targeted genomic locus. The presence of three distinct INDELs in clone 3B8, and two INDELs in an approximate 2:1 ratio in clones 3C12 and 3J7, is consistent with the known trisomy of chromosome 8 in HeLa cells. **B)** Representative Western blot image of HeLa WT cells (lane 1), HeLa VPS13B KO cells (clone 2D9) (lane 2) and VPS13B KO cells transiently transfected with a VPS13B expressing plasmid (lane 3).

**Figure S2.**
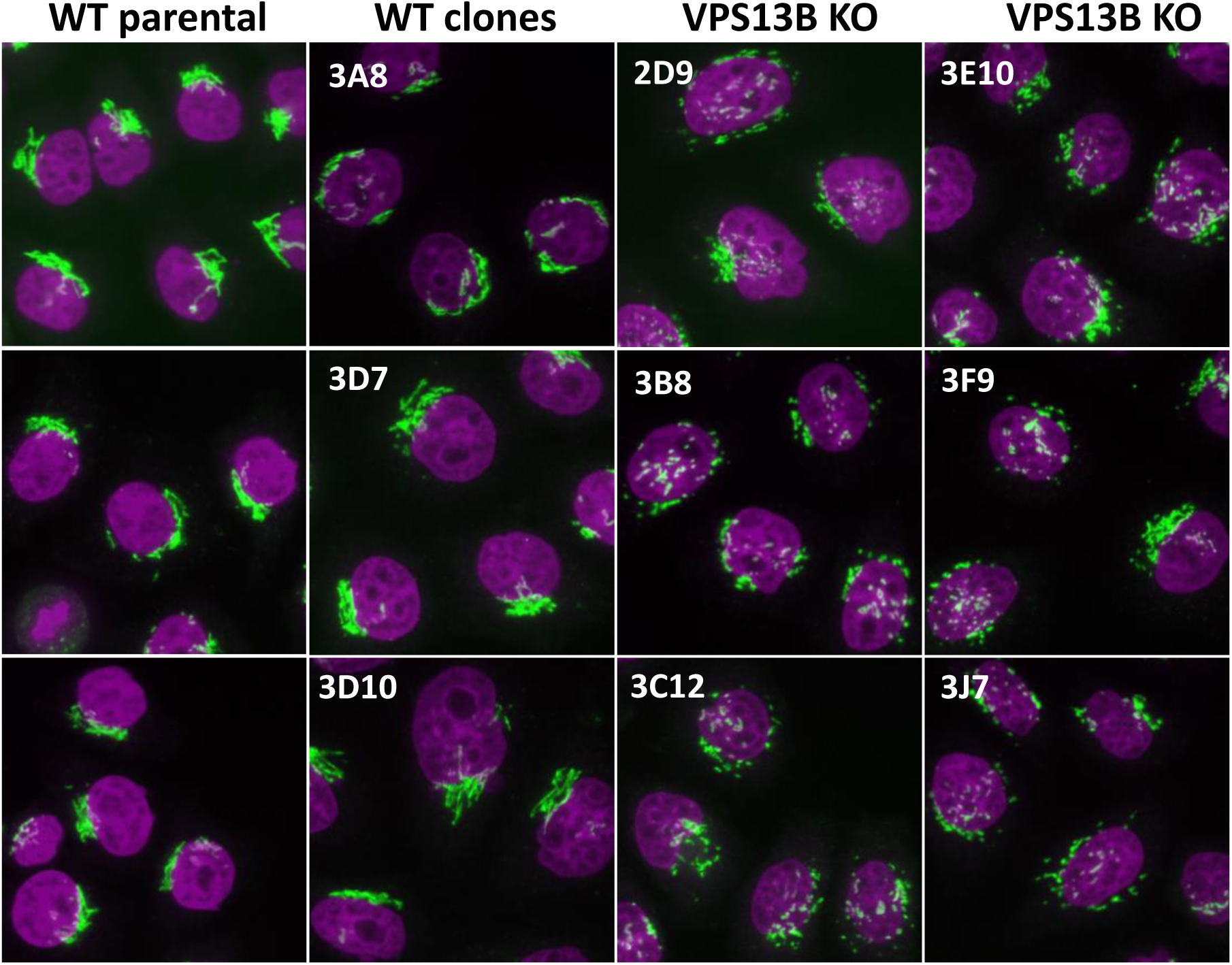
Microscopy images of VPS13B KO clones. Representative images of GM130-stained parental wild-type (WT) HeLa cells and VPS13B knockout (KO) or WT clones, illustrating a consistent genotype–phenotype correlation across all analyzed clones. Nuclei are counterstained with Hoechst 33342 (magenta).

**Figure S3.**
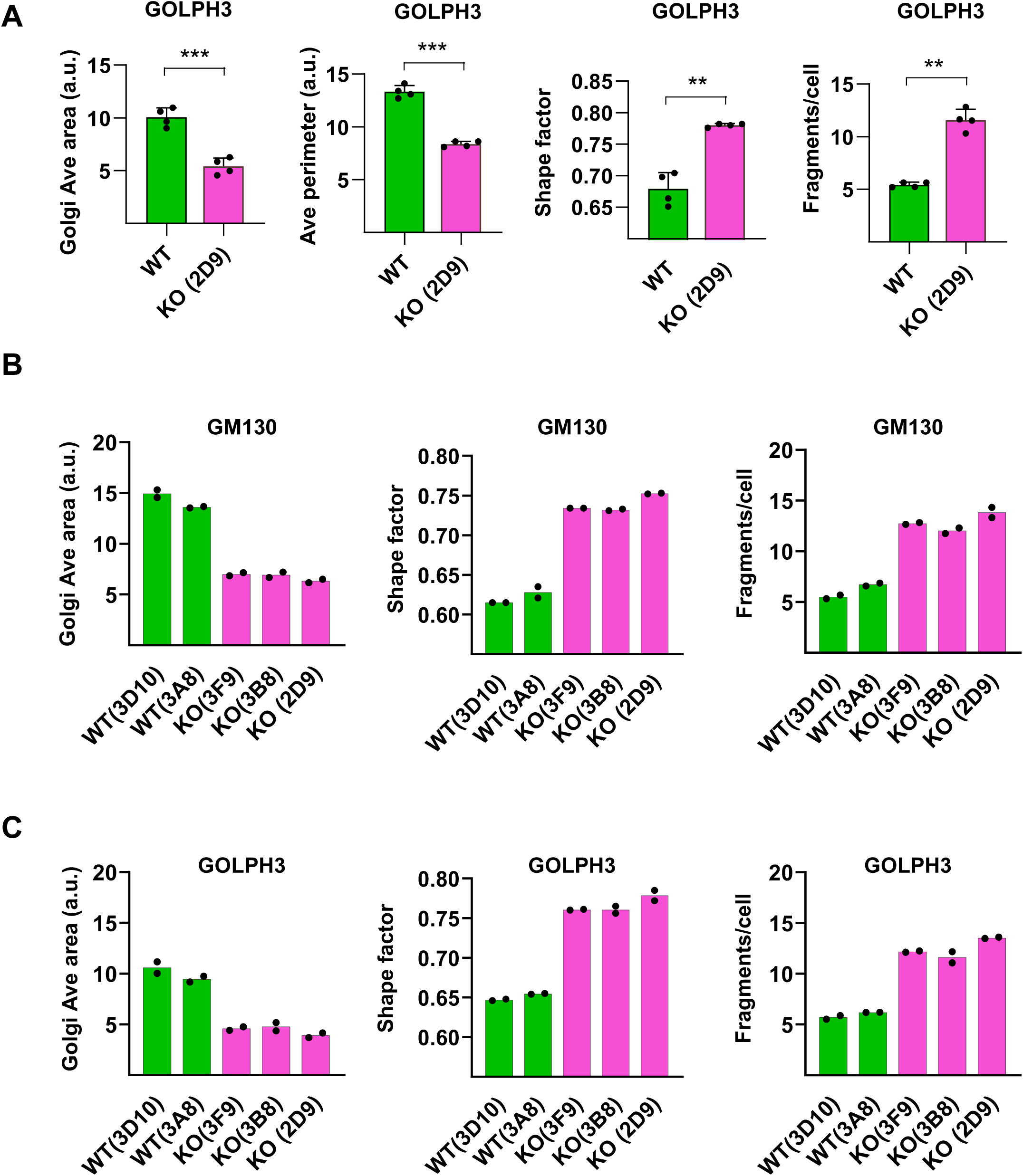
High-content image analysis of Golgi morphology in individual clones. **A)** Analysis of Golgi morphology parameters extracted from GOLPH3 images in WT and VPS13B KO (clone 2D9) HeLa cells. Average area and perimeter of Golgi fragments (Area Ave, Perimeter Ave); Shape factor average; average number of Golgi fragments per cell. Analysis based on 1000-1500 cells/point (**p<0.01; ***p<0.001; paired t-test, 2-tails; N=4). **B, C)** Analysis of Golgi morphology parameters extracted from GM130 **(B)** and GOLPH3 **(C)** images in WT (clones 3D10 and 3A8) and VPS13B KO (clones 3F9, 3B8 and 2D9) HeLa cells. Average area of Golgi fragments (Area Ave, Perimeter Ave); Shape factor average; average number of Golgi fragments per cell. Analysis based on 1000-1500 cells/point.

**Figure S4.**
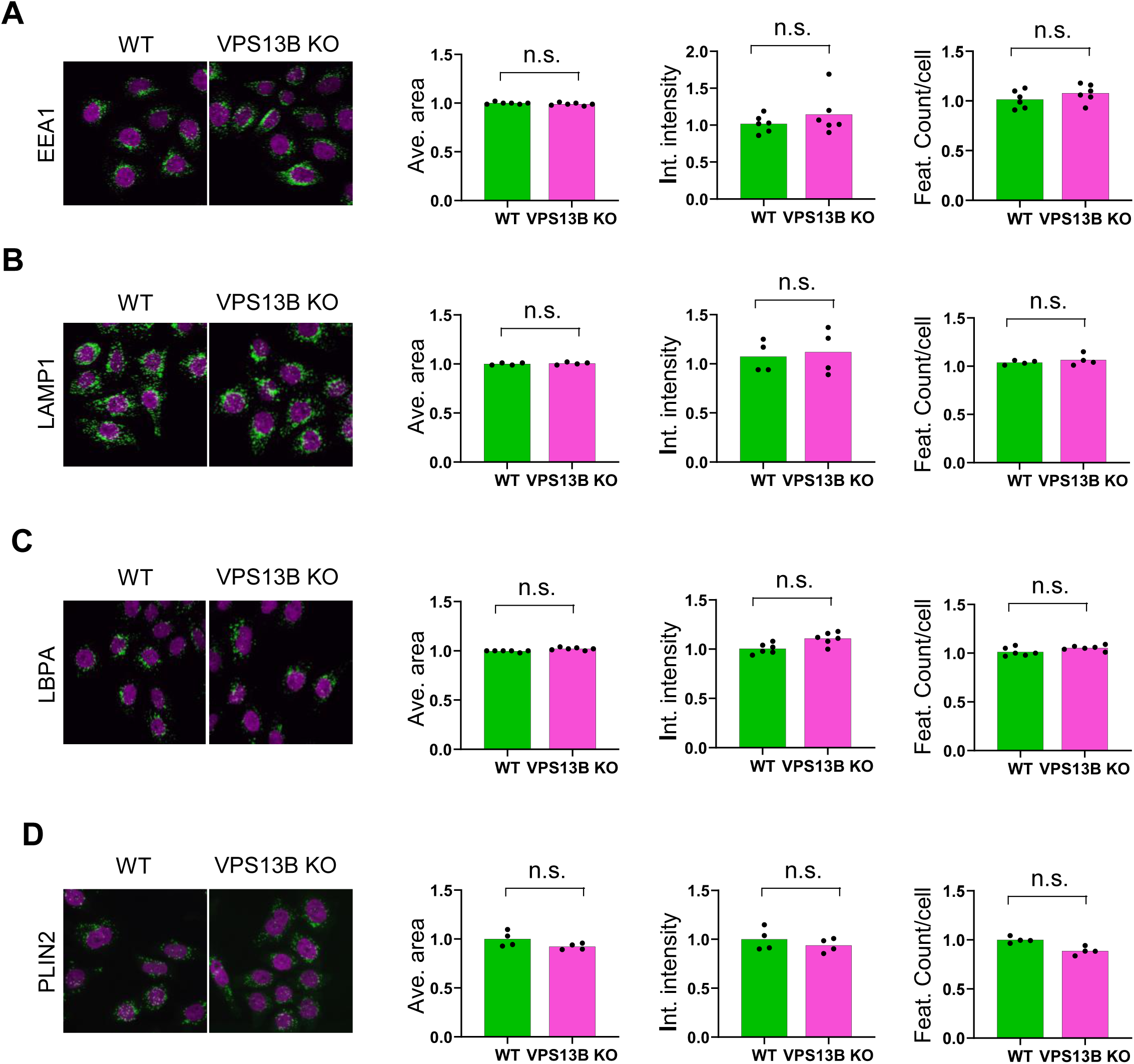
Endo/lysosomal and lipid droplet staining in VPS13B KO cells. Representative images and image analysis of WT and VPS13B KO (clone 2D9) HeLa cells stained with EEA1 **(A)**, LAMP1 **(B)**, LBPA **(C)** and PLIN2 **(D)** markers. Analysis displays values for Object average area, integrated signal intensity per cell, features count per cell. Analysis based on 1000-1500 cells/point. (paired t-test, 2-tails; N=4 (LAMP1, PLIN2), N=6 (EEA1, LBPA).

**Figure S5.**
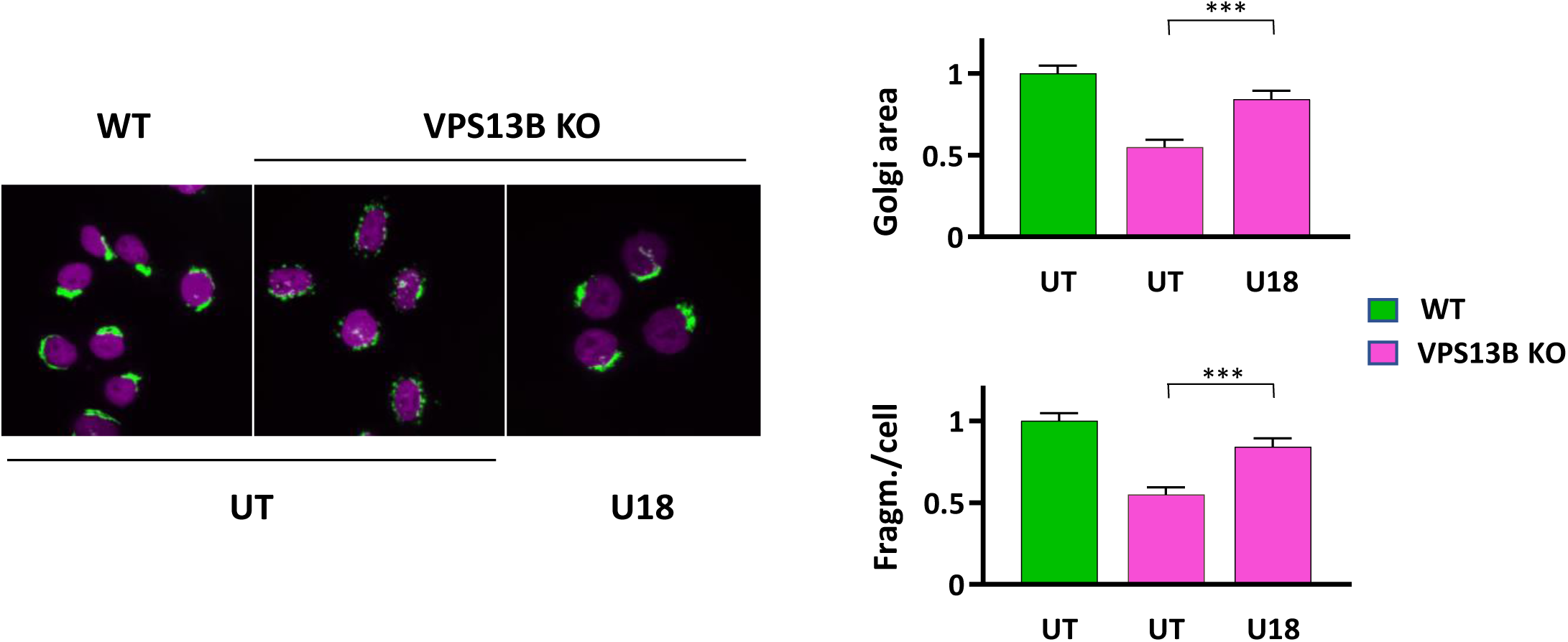
U18666A treatment recovers Golgi morphology in VPS13K KO cells. Representative images of WT or VPS13B KO treated or not with U18666A (U18) 10 µM for 24 hours. Cells were stained for GM130 (green) and Hoechst 33342 (magenta). Analysis of Golgi average area and average fragments per cell (values relative to WT UT cells). Analysis based on 1000-1500 cells/point. (***q<0.001. one-way ANOVA with Šidák correction; N=3)

**Figure S6.**
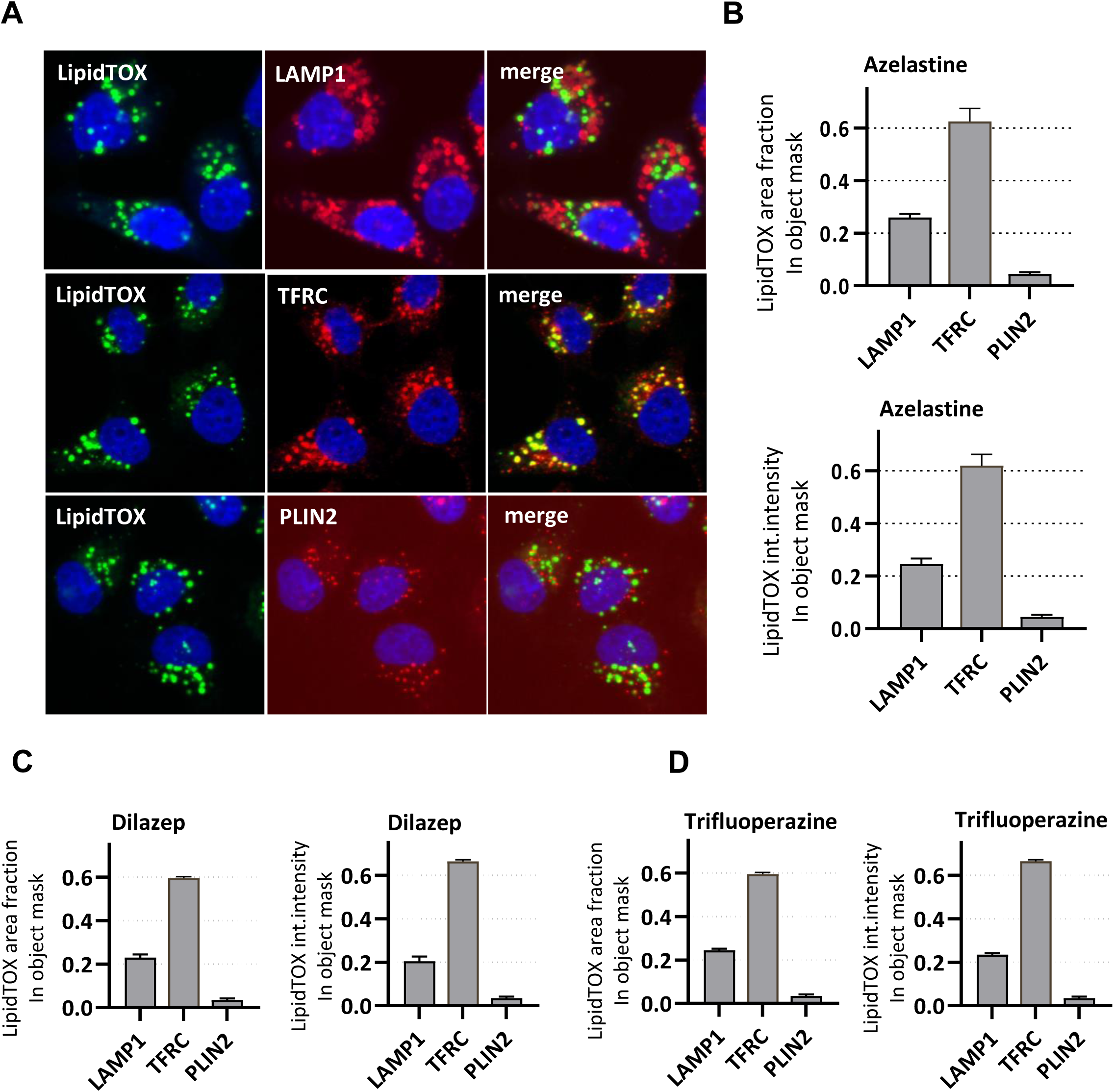
LipidTox subcellular localization in CAD-treated cells. **A)** Representative single-channel and merged confocal images showing co-staining of LipidTOX Green dye with markers of late endosomes/lysosomes (LAMP1), recycling endosomes (TFRC), and lipid droplets (PLIN2) in VPS13B knockout HeLa cells (clone 2D9) treated with azelastine for 24 hours. Cells were stained with the indicated markers, and images were acquired using an automated confocal microscope. **B)** Co-localization analysis was performed using MetaXpress® software by generating object masks for each marker channel. Quantification includes the fraction of LipidTOX-positive area (upper panel) or integrated LipidTOX signal intensity (lower panel) within LAMP1, TFRC, and PLIN2 object masks on azelastine-treated cells. Analysis based on 1000-1500 cells/point (N=2). **C, D)** Co-localization analysis performed as in panel B on dilazep-**(C)** and trifluoperazine-treated cells. Analysis based on 1000-1500 cells/point (N=2).

**Figure S7.**
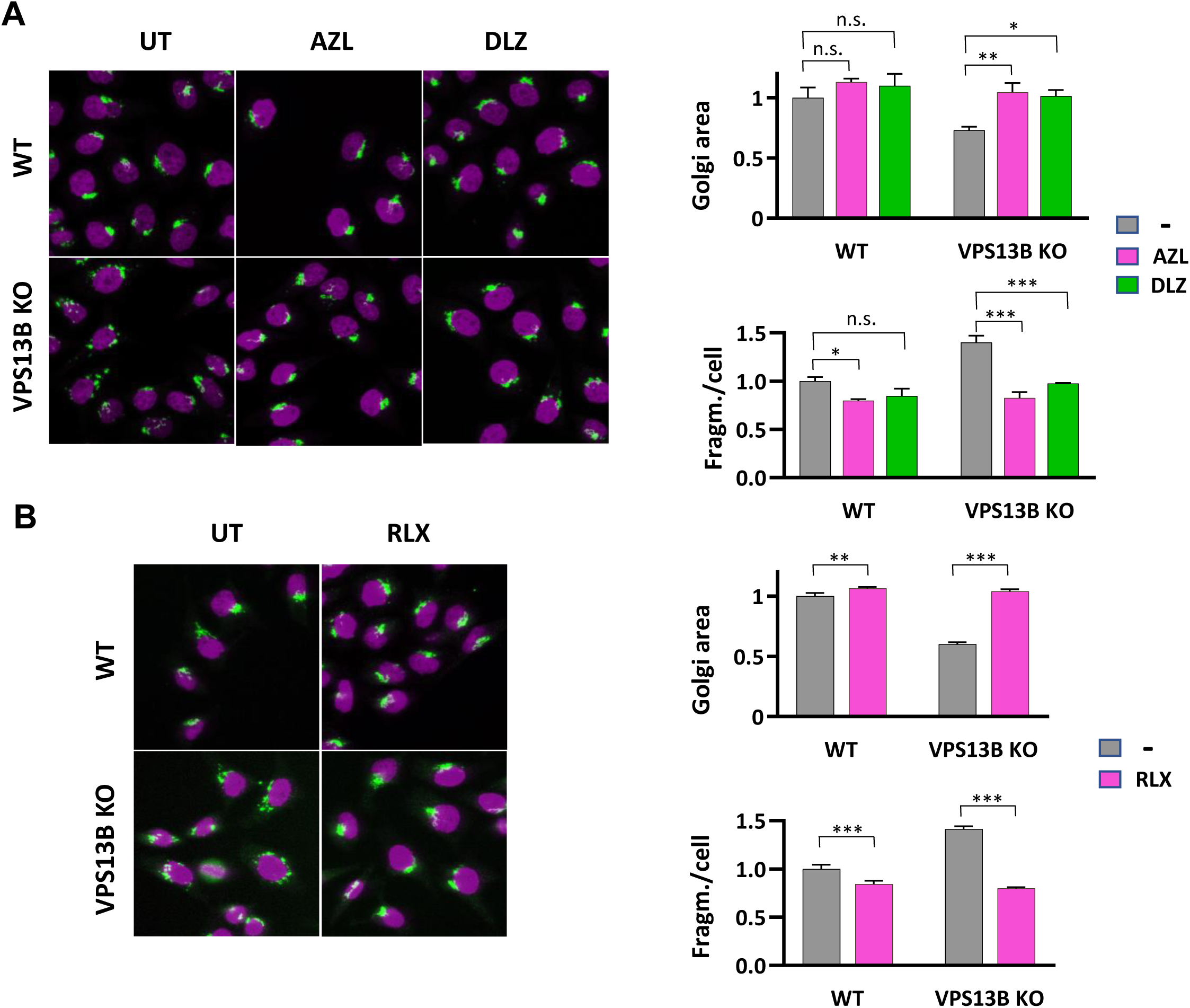
CAD effect on WT cells. Representative images of WT and VPS13B KO HeLa, either untreated (UT) or treated with azelastine (AZL) or dilazep (DLZ) 10 µM for 24 hours **(A)** or raloxifene (RLX) 10 µM for 24 hours **(B).** Cells are stained for GM130 (green). Nuclei are counterstained with Hoechst 33342 (magenta). Graphs illustrate Golgi average area and average fragments/cell (values relative to UT cells) quantified on 1000-1500 cells/point. (*q<0.05,**q<0.01,***q<0.001. two-way ANOVA with Tukey correction; N=3).

**Figure S8.**
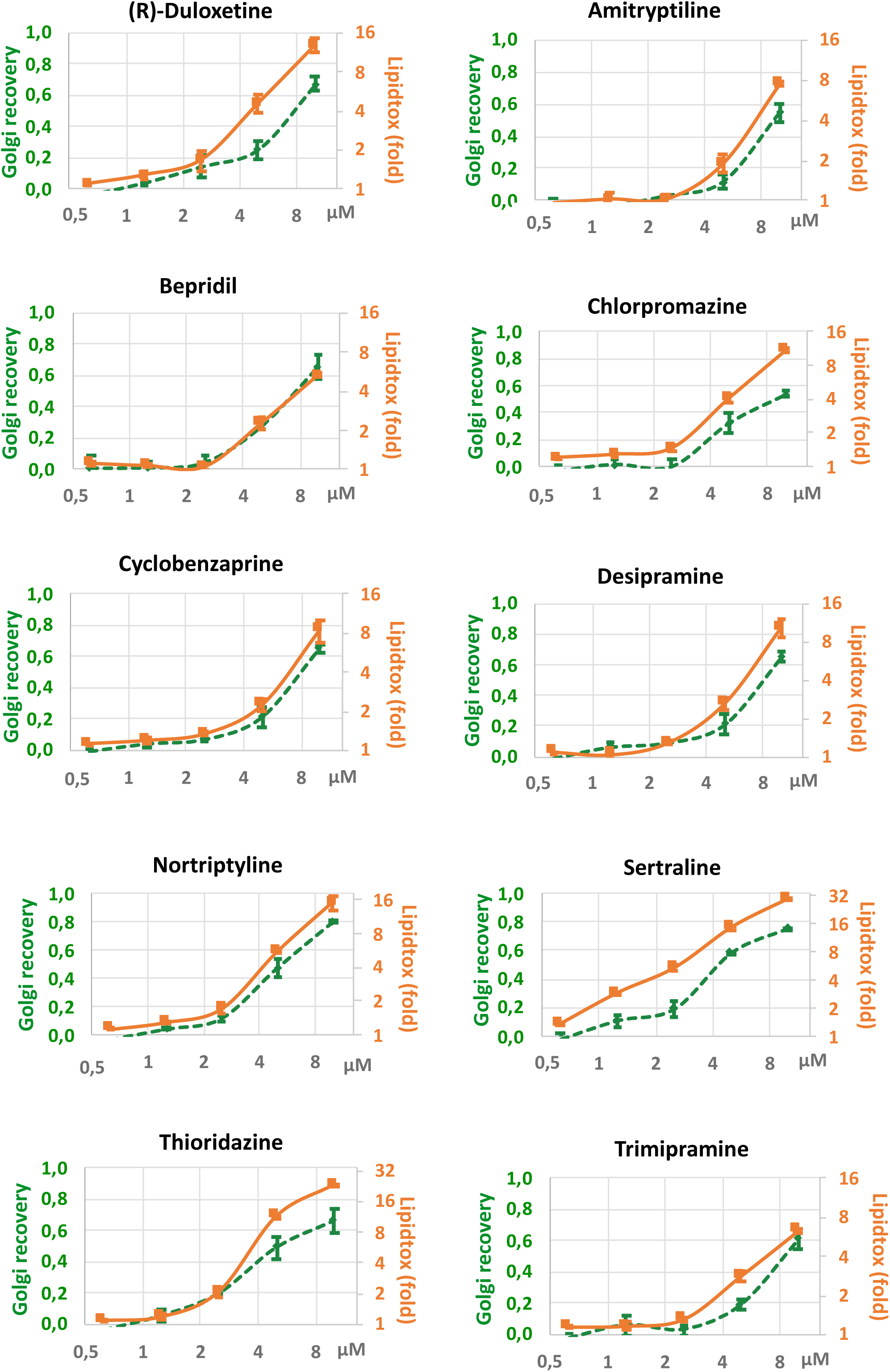
Dose-response curves of selected hit compounds. Dose-response curves of the indicated compounds displaying the Golgi recovery, expressed as the inverse (1-D) of 4-parametric distance to WT (green line) and LipidTox integrated intensity (relative to WT, a.u., orange line) Analysis based on 1000-1500 cells/point (N=2).

**Figure S9.**
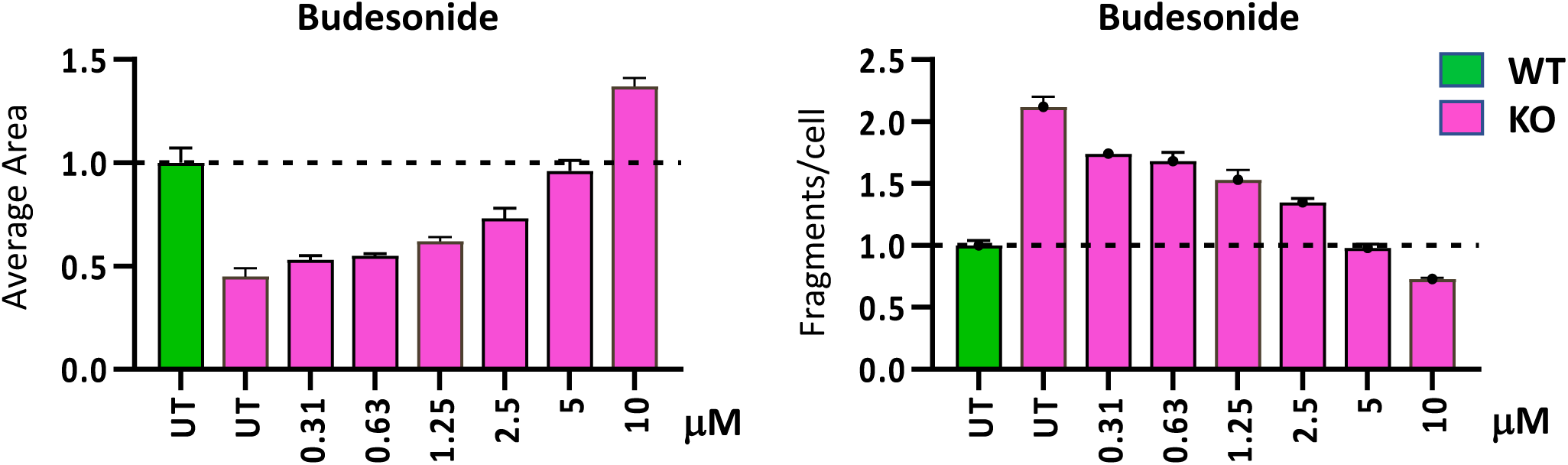
Area and fragment analysis in budesonide-treated cells. .Concentration-dependent variation of Golgi fragment Average Area and fragment count per cell from the dose response curve of budesonide (same data were used in figure 4D to calculate Golgi recovery) (N=2).

**Figure S10.**
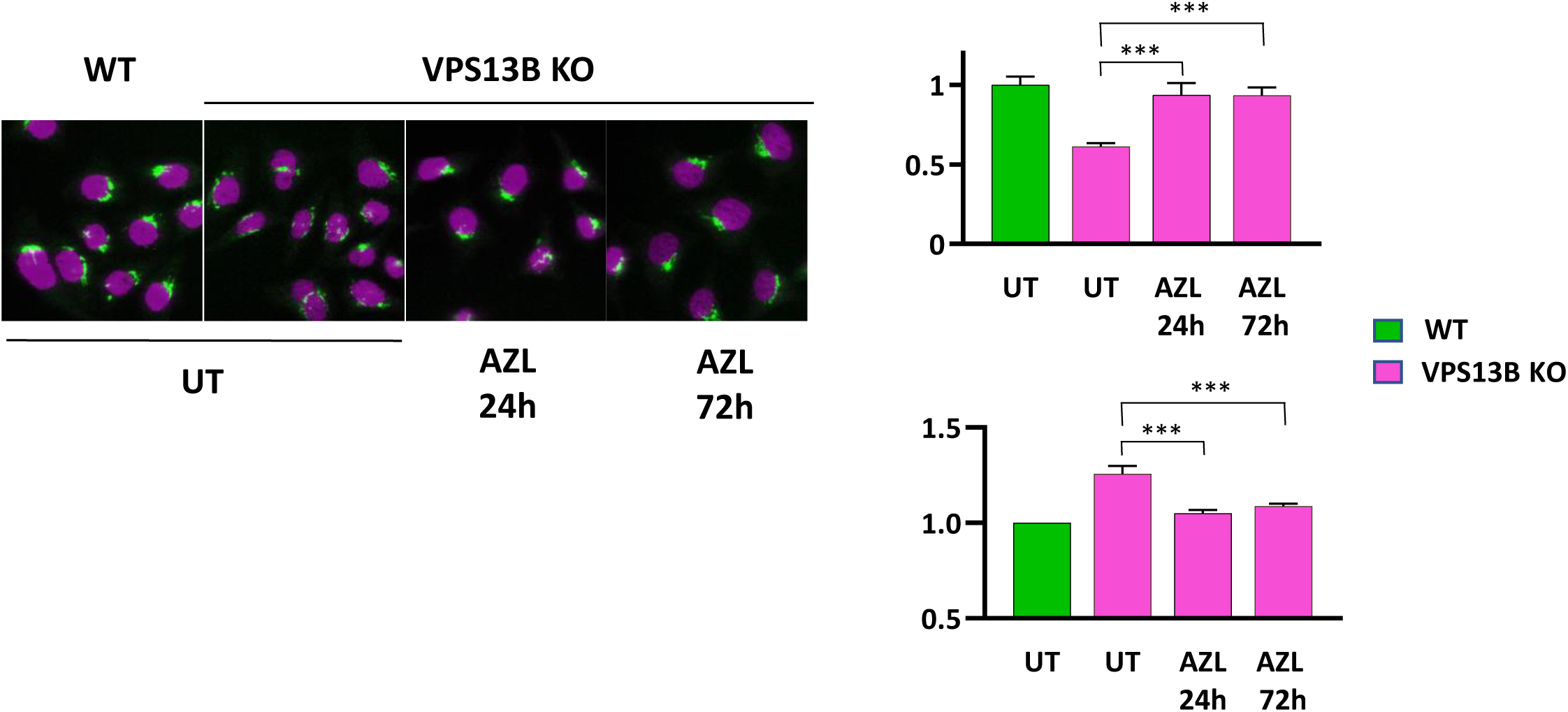
**Prolonged azelastine effect on Golgi recovery**. Representative images of WT and VPS13B KO HeLa, either untreated (UT) or treated with azelastine (AZL) 10 µM for 24 or 72 hours. Cells are stained for GM130 (green). Nuclei are counterstained with Hoechst 33342 (magenta). Graphs illustrate Golgi average area and average fragments/cell (values relative to UT cells) quantified on 1000-1500 cells/point (***q<0.001. one-way ANOVA with Šidák correction; N=3).

**Figure S11.**
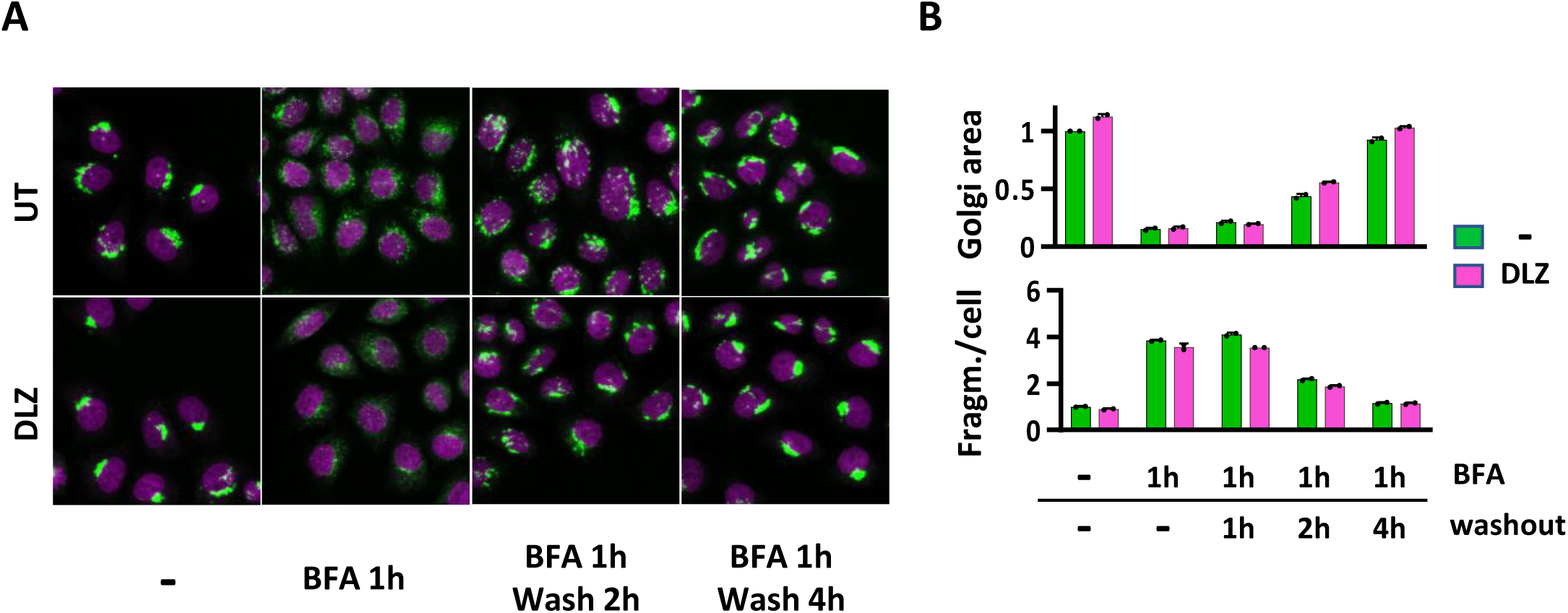
Dilazep do not prevent brefeldin A-induced Golgi fragmentation. **A)** Representative images of WT HeLa cells treated (DLZ) or not (UT) with dilazep 10 µM for 24 hours. Where indicated, cells have been subsequently with brefeldin A (BFA) 1 µg/ml for 1 hours and after 3 washes, let recover in full medium for 2 or 4 additional hours. In DLZ samples, the compound has been kept in all the course of the experiment. Cells are stained for GM130 (green). Nuclei are counterstained with Hoechst 33342 (magenta). **B)** Graphs illustrating Golgi average area and average fragments/cell (values relative to UT cells) quantified on 1000-1500 cells/point (N=2).

**Figure S12.**
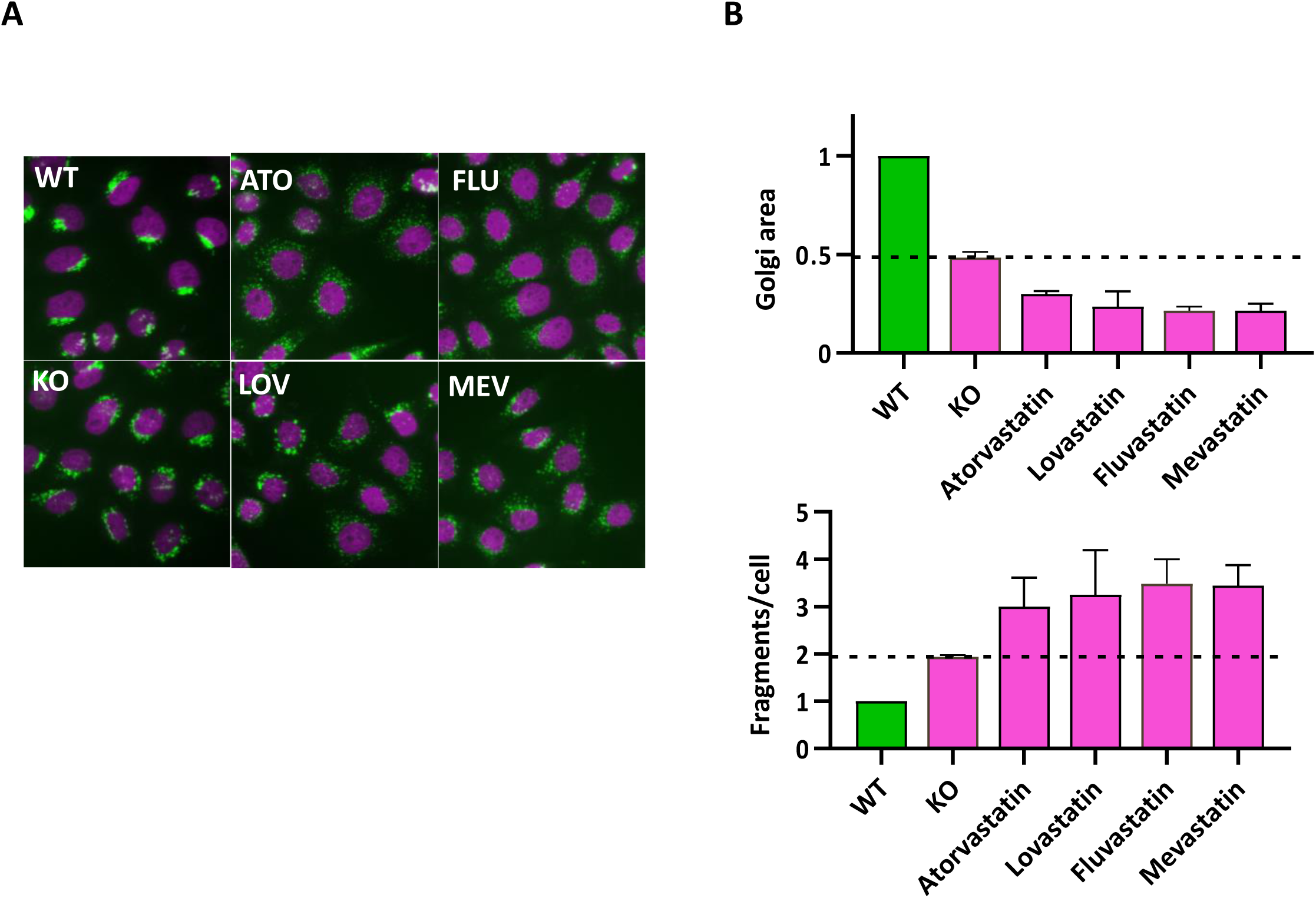
Statins induce Golgi fragmentation. **A)** Representative images from the Prestwick chemical library screening illustrating GM130 staining (green) WT and VPS13KO HeLa cells untreated (KO) or treated with the indicated compounds at 10 µM for 24 hours: Atorvastatin (ATO); lovastatin (LOV); Fluvastatin (FLU); Mevastatin (MEV). Nuclei are stained with Hoechst 33342 (magenta) **B)** Graphs illustrating Golgi fragment average area and average Golgi fragments/cell (values relative to UT cells) quantified on 1000-1500 cells/point (N=2).

**Figure S13.**
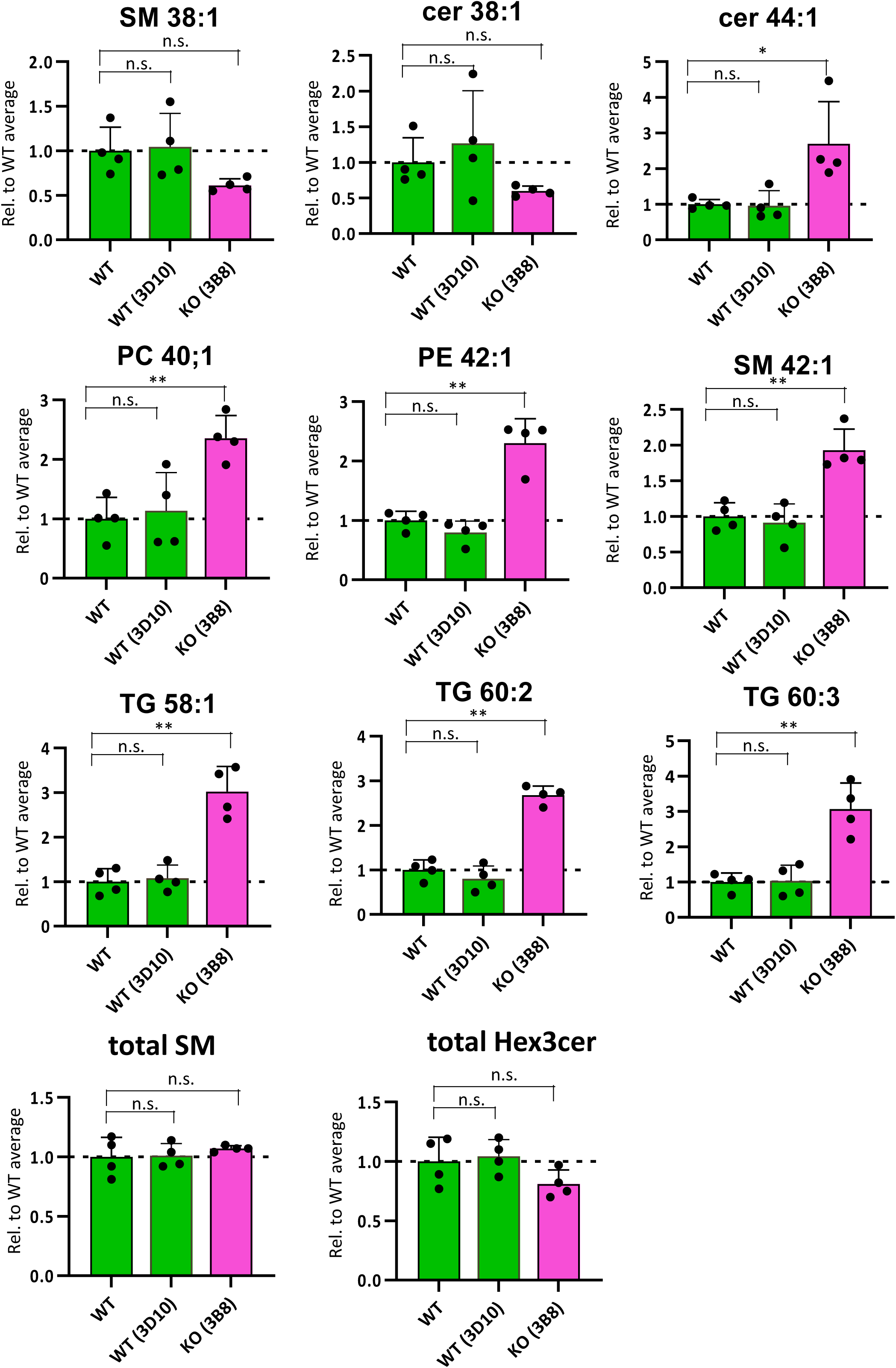
Lipid species altered in VPS13B KO clones. Relative amounts of the indicated lipid species (calculated as a percentage of the respective class) in WT parental cells (WT), in WT cells (clone 3D10) and in VPS13B KO cells (clone 3B8). (*p<0.05, **p<0.01. one-way ANOVA with Dunnett correction; N=4).

**Figure S14.**
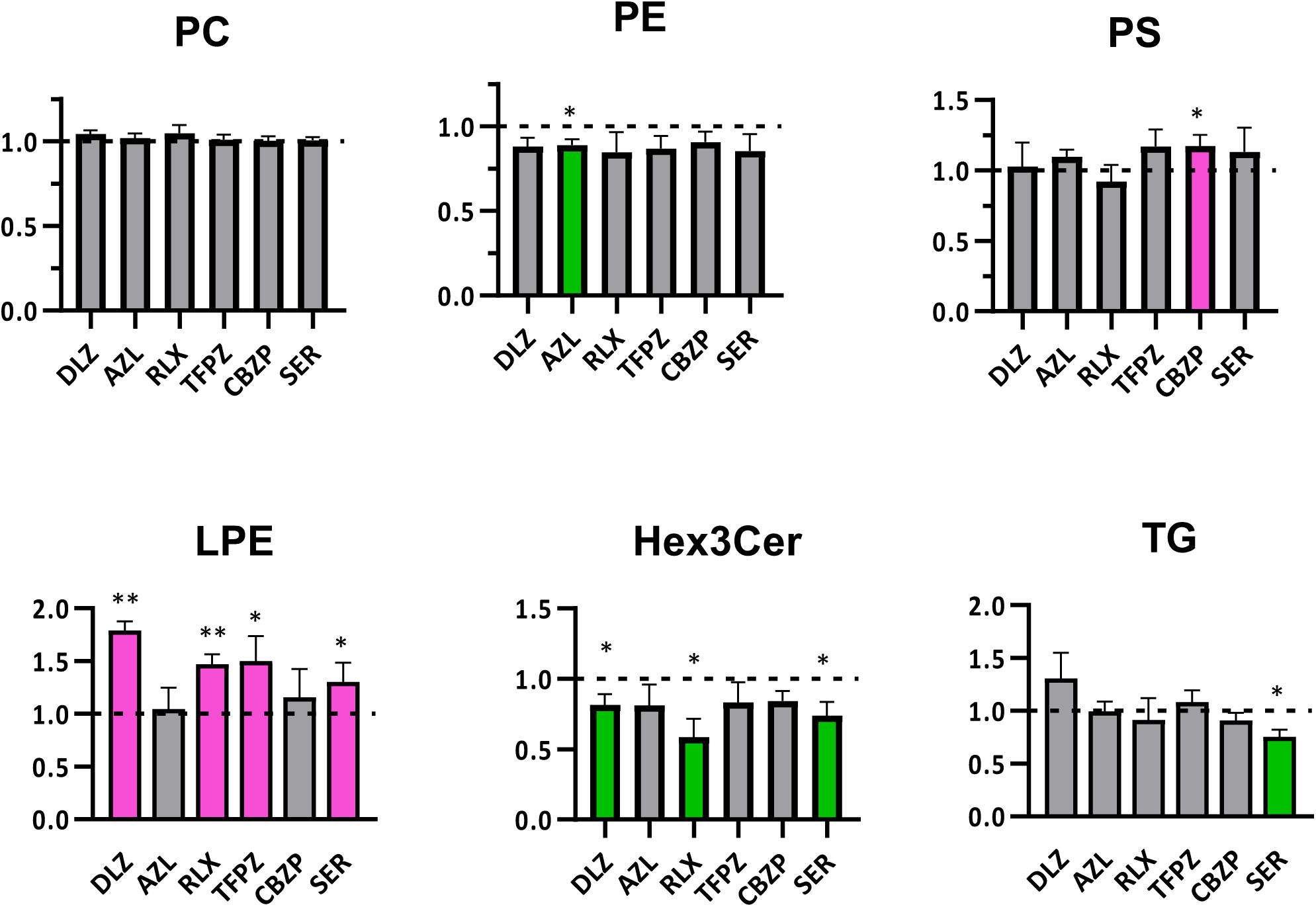
Lipid classes alterations induced by drug treatments. Abundance of indicated lipid classes in VPS13B KO cells (clone 2D9) treated with the indicated compounds (10 µM, 24 hours), relative to untreated cells. DLZ (dilazep); AZL (azelastine); RLX (raloxifene); TFPZ (trifluoperazine); CBZP (cyclobenzaprine); SER (sertraline). (*q<0.05, **q<0.01. multiple paired t-tests with Benjamini-Hochberg correction; N=4).

**Figure S15.**
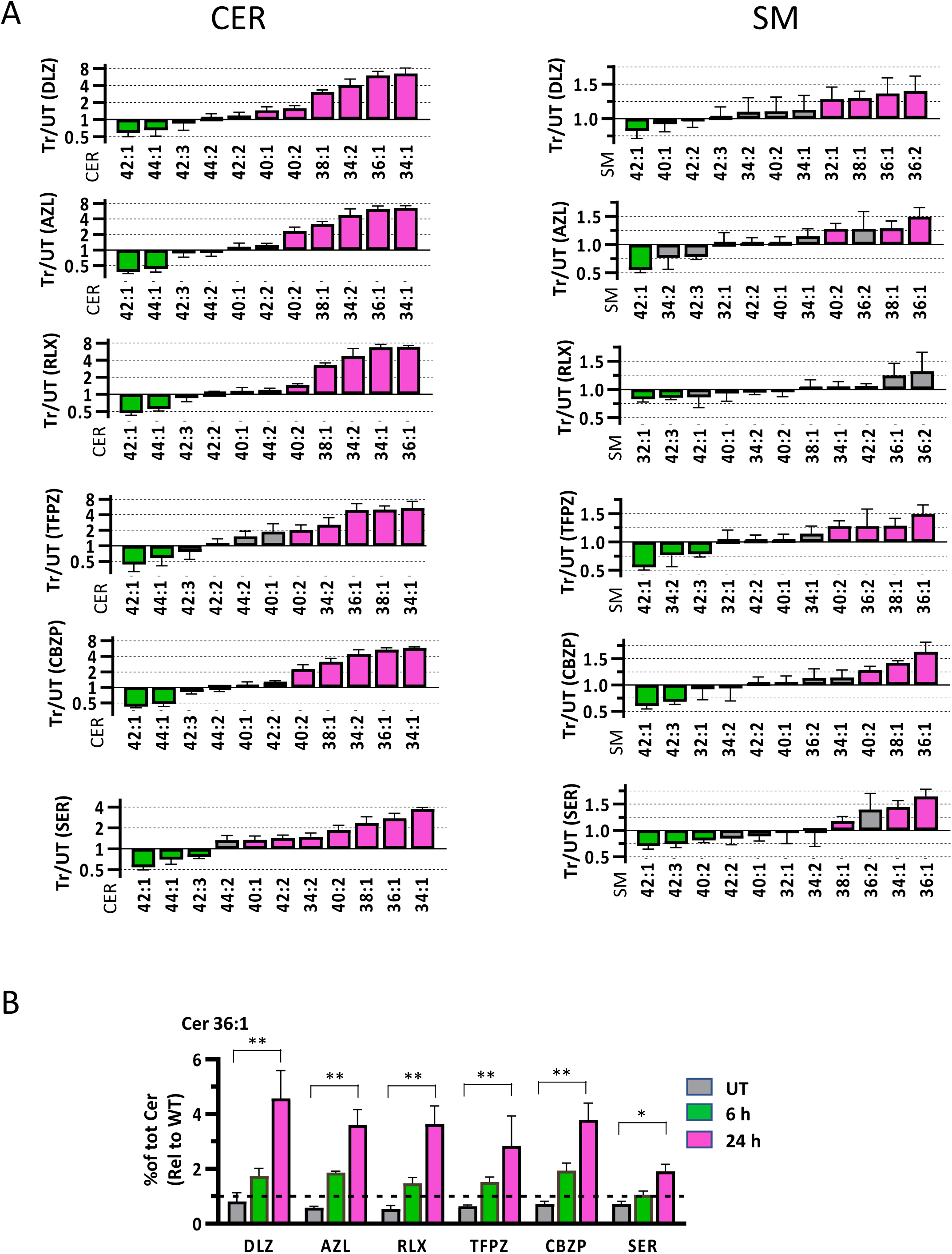
Relative ceramide and sphingomyelin alterations in drug-treated cells. **A)** Ratios of ceramide (CER) and sphingomyelin (SM) lipid species in treated (Tr) versus untreated (UT) cells. Values for individual lipid species were normalized to their respective lipid class prior to ratio calculation. Cells were treated for 24 hours at 10 µM with the following compounds: DLZ (dilazep), AZL (azelastine), RLX (raloxifene), TFPZ (trifluoperazine), CBZP (cyclobenzaprine), and SER (sertraline). Color coding indicates statistical significance: magenta for ratios >1 and green for ratios <1, both with p < 0.05 (multiple paired ratio t-tests, N = 4). Non-significant changes (ratios not different from 1) are shown in grey. **B)** Relative amounts of CER 36:1 (calculated as a fraction of total SM) in VPS12B KO cells treated with the indicated compounds for 6 or 24 hours. (*p<0.05, **p<0.0001. one-way ANOVA with Šidák correction; N=4).

**Figure S16.**
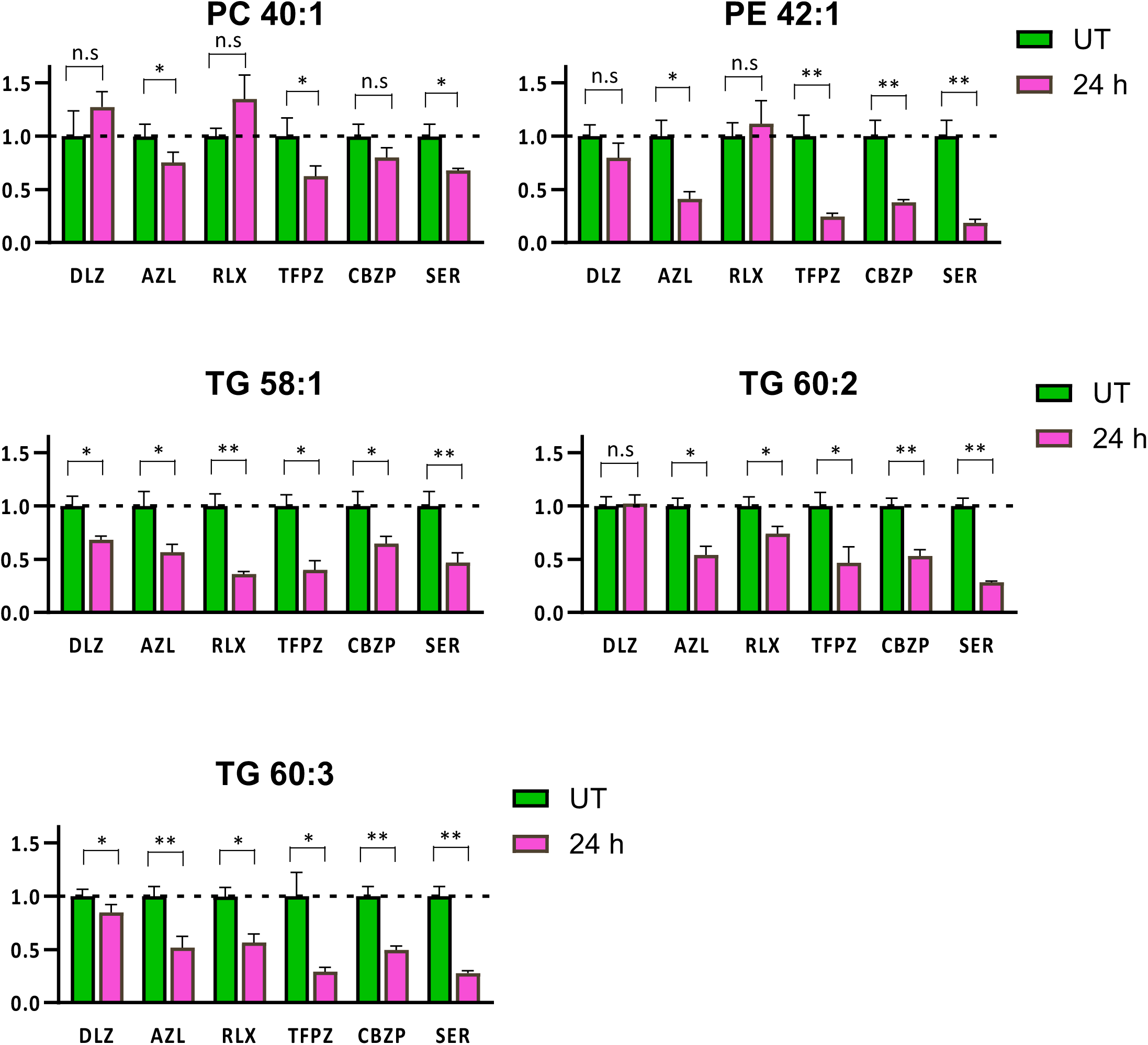
Drug-induced alteration of selected phosphatidylcholine (PC), Phosphatidylethanolamine (PE) and triacylglycerol (TG) species. Relative amounts of the indicated lipid species in VPS12B KO cells treated with the indicated compounds for 24 hours. (*p<0.05, **p<0.01. multiple paired ratio t-tests with Benjamini-Hochberg correction; N=4).

**Figure S17.**
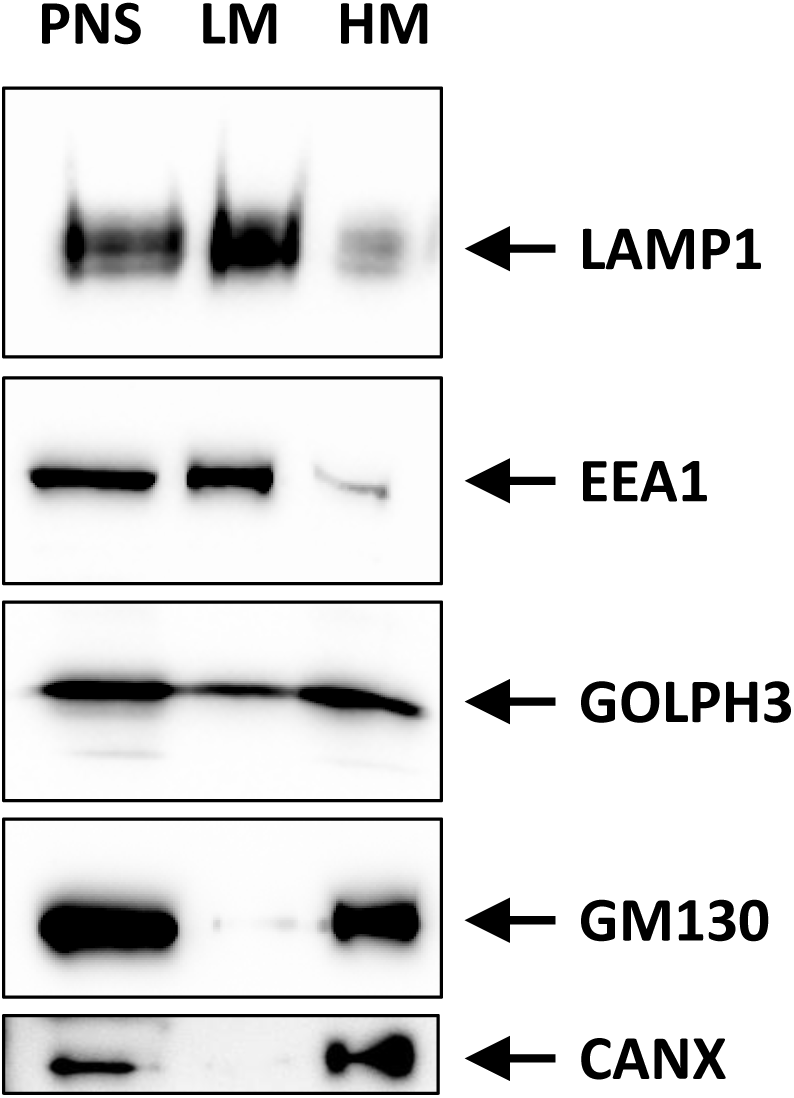
**Representative Western blot images of subcellular fractions from HeLa cells**: post-nuclear supernatant (PNS), light membranes (LM), heavy membranes (HM) probed with the indicated antibodies.

**Figure S18.**
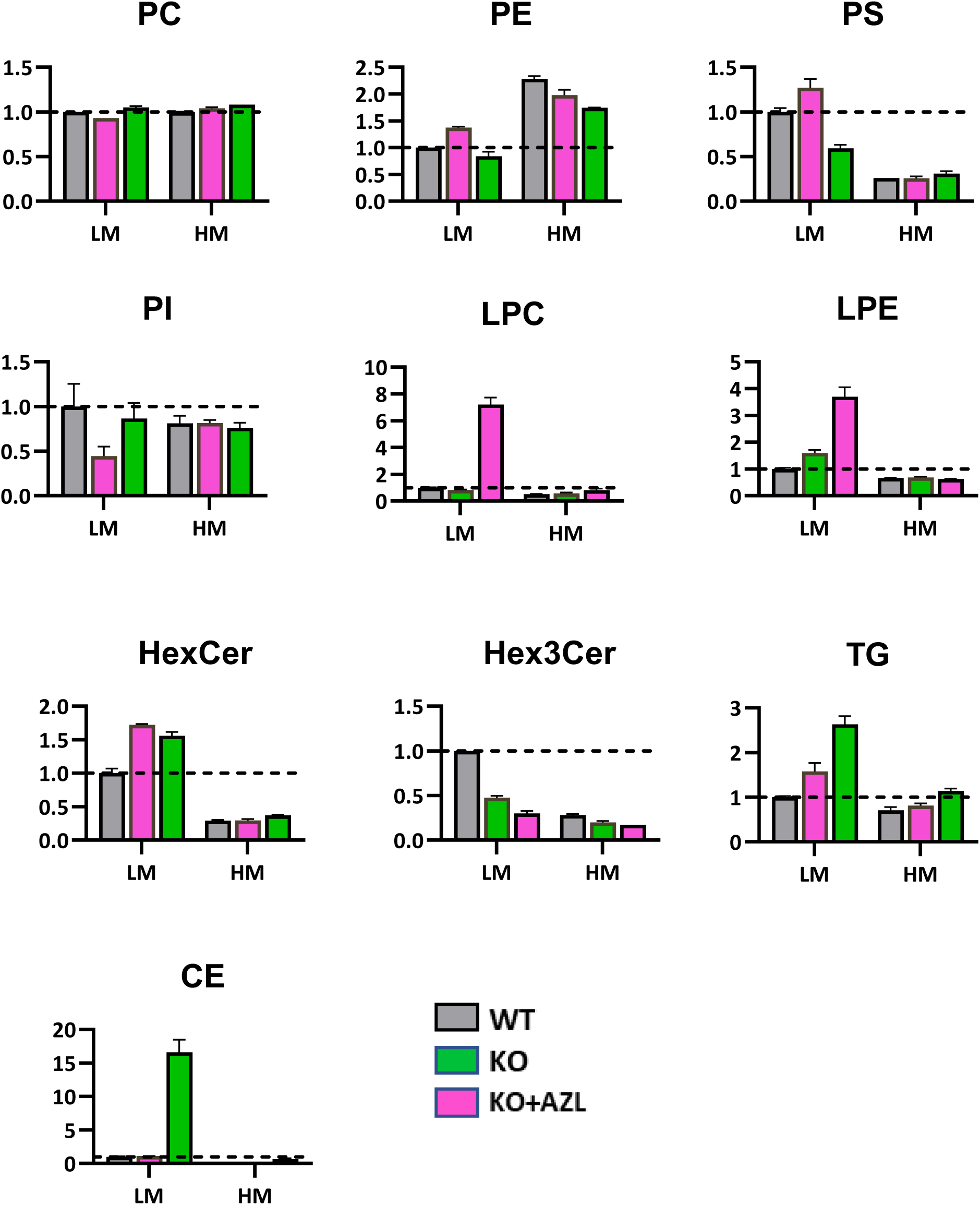
**Relative lipid classes abundance from LM and HM fractions In AZL-treated cells**. Relative amounts of total PC, PE, PS, PI, LPC, LPE, HexCer, HexCer, TG, CE in light (LM) or Heavy (HM) membranes prepared from VPS12B KO cells treated (AZL) with not (UT) with TFPZ 10 µM for 24 hours (N=2). Data are normalized to the LM value in UT cells.

**Figure S19.**
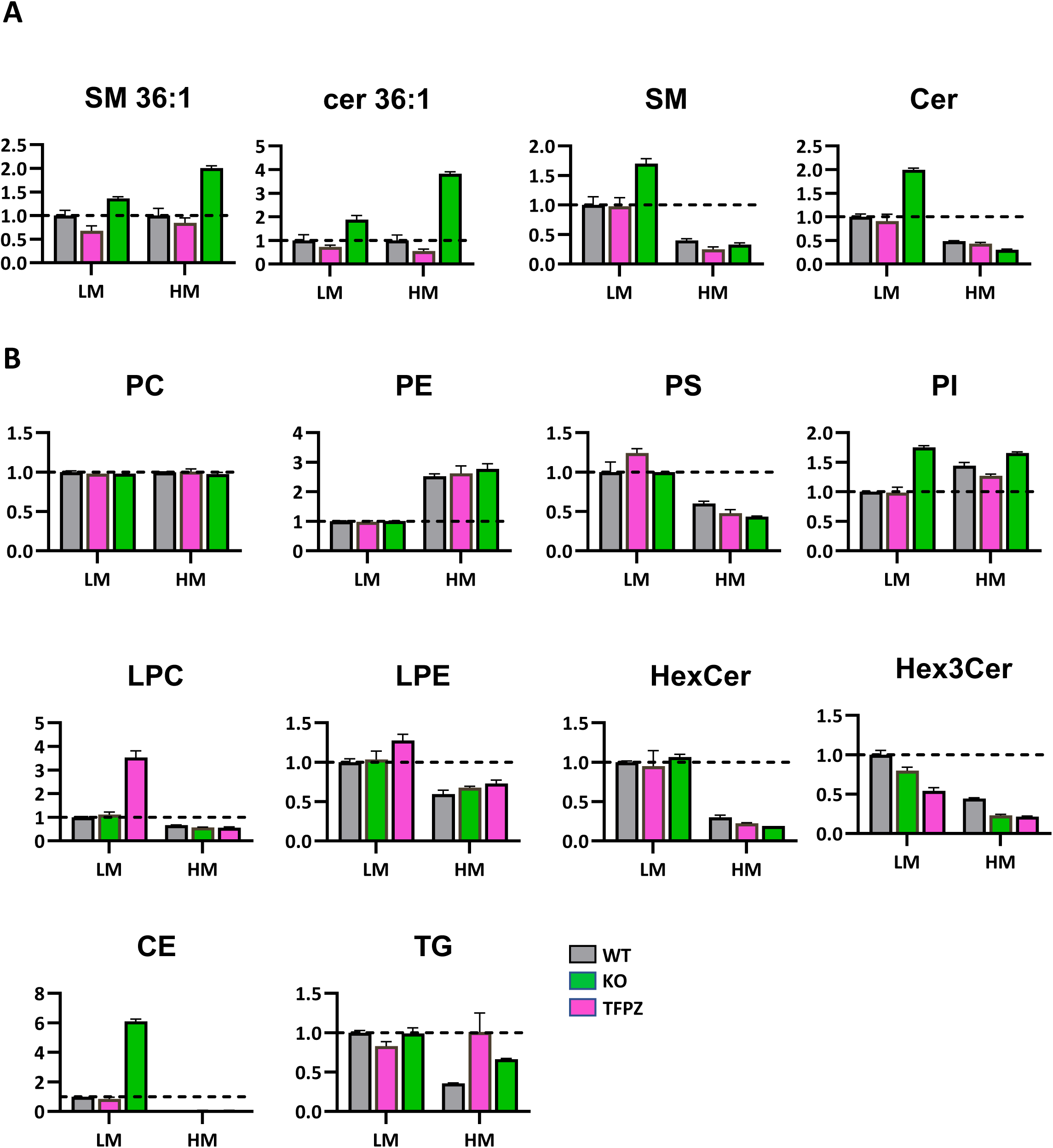
**Relative lipid abundance from LM and HM fractions In TFPZ-treated cells**. D) Relative amounts of total SM, total CER, SM 36:1 (calculated as a fraction of total SM) and CER 36:1 (calculated as a fraction of total CER) in light (LM) or Heavy (HM) membranes prepared from WT and VPS12B KO cells treated (TFPZ) with not (UT) with TFPZ 10 µM for 24 hours. **B)** Relative amounts of total PC, PE, PS, PI, LPC, LPE, HexCer, HexCer, TG, CE in light (LM) or Heavy (HM) membranes prepared from WT or VPS12B KO cells treated (TFPZ) with not (UT) with TFPZ 10 µM for 24 hours (N=2). Data are normalized to the LM value in UT cells.

**Figure S20.**
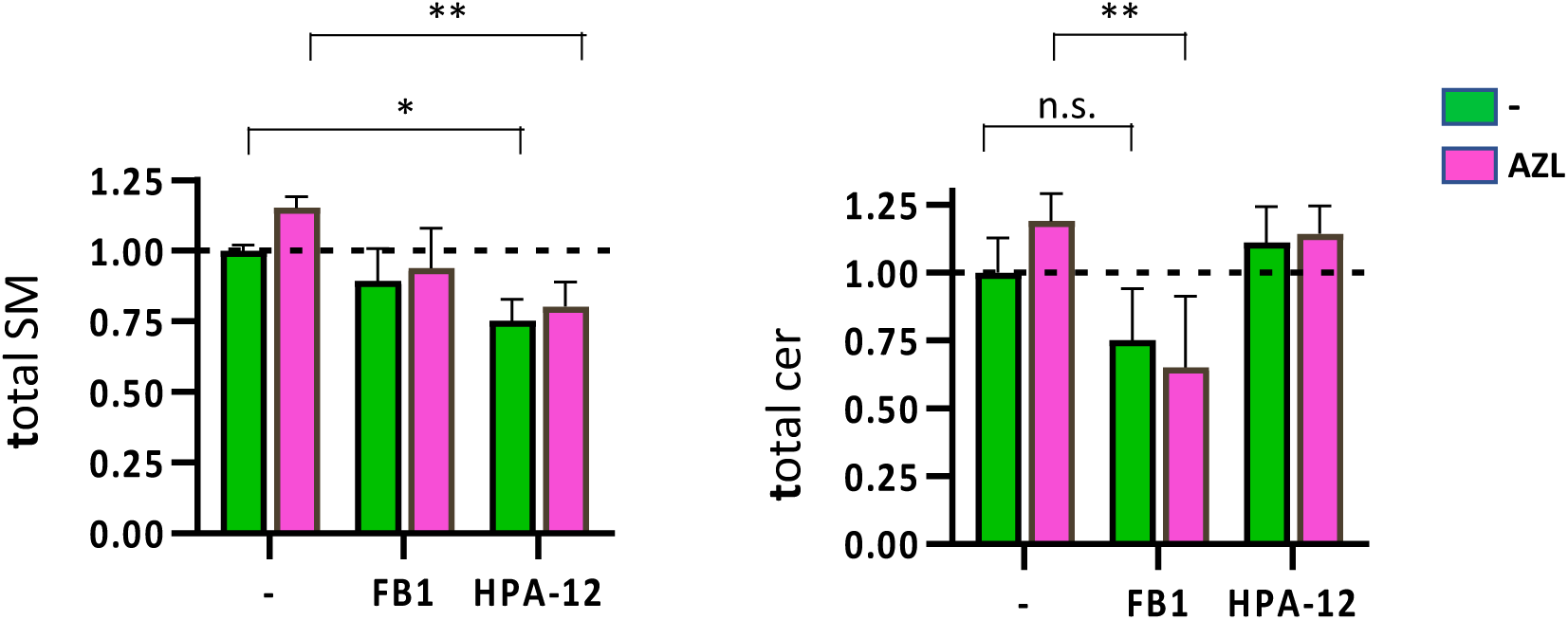
Effect of sphingolipid synthesis inhibitors of Sphingomyelin and ceramide azelastine-induced changes. Relative amounts of total SM and total CER in cells treated (AZL) or not (UT) with 10µM AZL for 6 hours in combination with 20 µM fumonisin B1 (FB1) or 5 µM HPA-12. (*p<0.05, **p<0.01. one-way ANOVA with Šidák correction (N=3).

**Figure S21.**
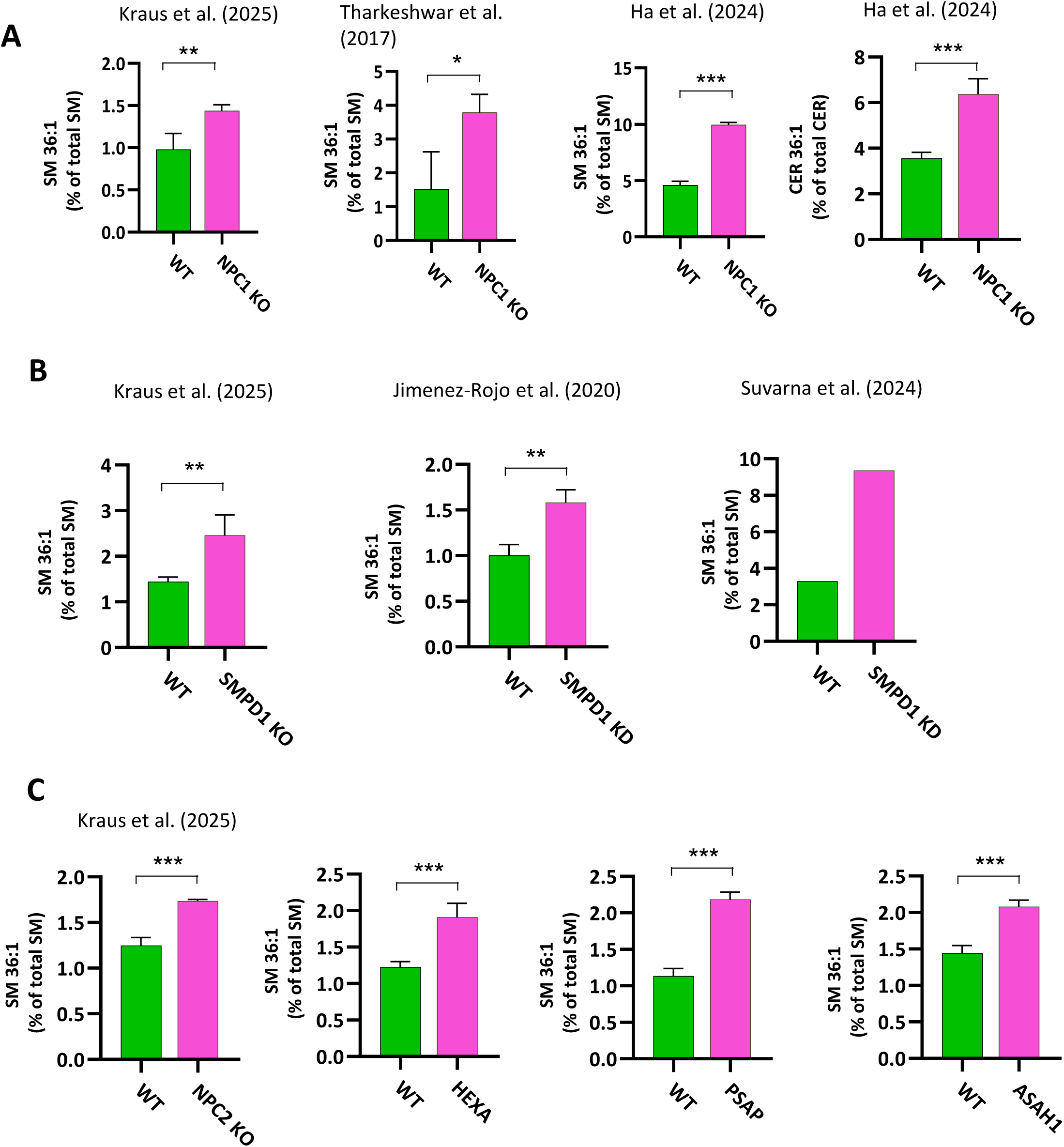
**SM 36:1 levels in re-analized lipidomic datasets**. Lipid content data obtained from published datasets were re-analyzed expressing the levels of SM 36:1 CER 36:1 as a percentage of total SM and CER content respectively, in cellular KO or KD models of NPC1 **(A)** SMPD1 **(B)** and other lysosomal storage disorder-causing genes **(C)**. Cell line used is HeLa, except Ha et al. (using HEK293) and Suvarna et al. (using A673). All references are cited in the main text. Full data are in Suppl. File X. (*p<0.05, **p<0.01, ***p<0.001. paired t-test, 2-tails)

**Figure S22.**
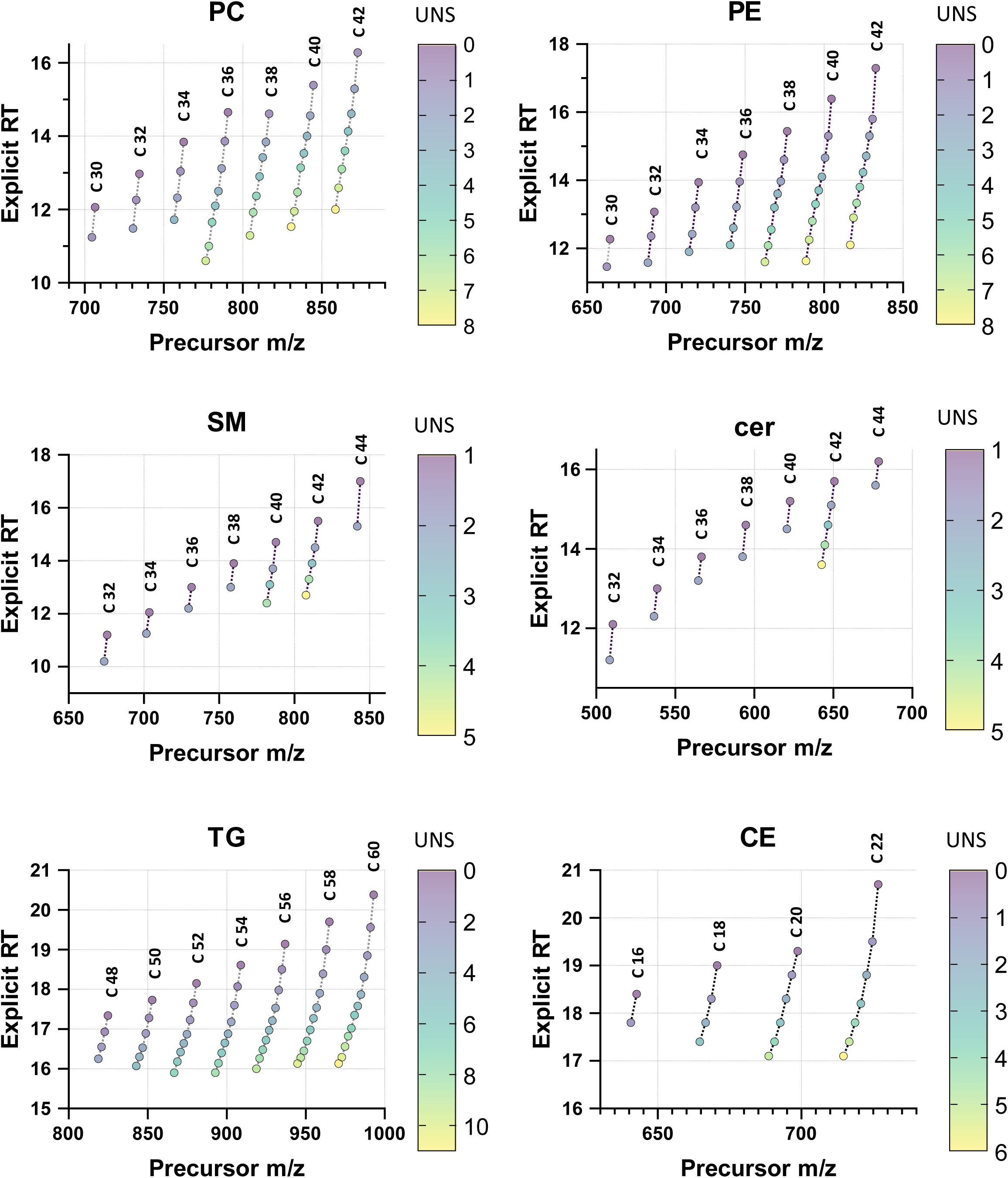
Precursor ion m/z-retention time relationship of analysed lipids. Graphs showing the relationship between precursor ion m/z and explicit retention time (min) for lipid species within the indicated lipid classes. Dotted lines connect lipid species with the same total number of acyl chain carbons. Colors indicate the total number of double bonds (degrees of unsaturation, UNS).

## Notes

### Competing Interest Statement

The authors have declared no competing interest.

### Summary of Updates

This version of the manuscript has been revised to respond to part of the Reviewer comments from the Review Commons platform. Further planned experiments are the detailed in the Revision Plan.

